# Longevity Bottlenecks

**DOI:** 10.1101/2023.08.18.553936

**Authors:** Michael Florea, Mark Hamalainen, Patrick Seebold, Nathan Cheng, Paul Murray, Alex James Colville, Sally Zheng, Dylan Ingham, Ridhi Kantelal, Rose De Sicilia, (Longevity Biotech Fellowship consortium)

## Abstract

The longevity field has received an influx of capital and talent over the past 5 years, but it is unclear where directing these resources would result in the biggest positive impact. We aimed to establish a systematic, rigorous and unbiased way to identify the areas where increased investment would accelerate progress across the whole longevity field the most. To do so, we surveyed ∼400 participants across various sectors of longevity, asking them to 1) identify the major bottlenecks they are experiencing, 2) list their most needed solution, and 3) rate the potential efficacy and barriers to development of various aging intervention strategies. We built a classification system of Bottlenecks and Solutions based on grouping related answers and found the most frequently listed bottlenecks to be 1) lack of validated aging biomarkers; 2) an overall lack of funding; and 3) lack of good models for aging studies. Surprisingly, the most wanted solution was greater availability of large public datasets. Indeed, a common theme across all answers was the need for a new data-centric structure of scientific research, where large datasets are routinely gathered and made available, access walls are removed, protocols are standardized, negative and unpublished data are shared, and AI systems are released on the data for automated discovery. Finally, a lack of regulatory clarity was listed as the biggest barrier to development across all interventions, whereas cellular reprogramming, organ replacement, and genetic medicine (gene therapies and gene editing) were perceived as the intervention strategies with the highest potential for increasing healthy lifespan. We provide these data as a resource for funding agencies, philanthropists, entrepreneurs and newcomers to the field as a means to identify high impact areas to fund and work on.

**Key takeaways:**

- 395 Participants were surveyed for their biggest bottlenecks and most needed solutions
- Top Bottlenecks: lack of Validated Biomarkers; Overall lack of Funding and Slow & Expensive Models.
- Top proposed Solutions: more Public Datasets; improved Regulatory Path; and Overall More Funding.
- Bottlenecks and Solutions vary substantially across industry areas.
- Rapamycin and calorie restriction are perceived as the most efficacious interventions in the near term.
- Somatic reprogramming, organ replacement, and genetic medicine are perceived as the most efficacious interventions in the long term (25 years).
- Sirtuin and NAD targeting therapies are seen as the least efficacious interventions in all time-frames.
- Across all interventions, Regulatory Issues are perceived as the most severely inhibiting factor in the development of the intervention.

## Introduction

The longevity field has seen tremendous growth in the past 5 years. Multiple highly capitalized and visible companies were founded and several new philanthropic and venture funds were launched between 2020-2023 (*1, 2*). This increased visibility and resource pool is a welcome development for a field long lacking resources. However, there is a lack of clarity or consensus on where investment and talent would make the biggest impact. In academia, funding is allocated based on the visibility, scientific credentials, and grantsmanship of individual investigators. In Industry, funding is allocated based on perceived revenue potential and fundraising ability of individual companies. Little systematic effort has been dedicated to identifying systemic deficiencies which retard or prevent progress for the field overall - to our knowledge, no such analysis has ever been published. As a result, the allocation of funding and attention is not coordinated and instead often proceeds as hype cycles where certain interventions are overfunded while other, critical areas and promising interventions receive little attention and remain underdeveloped.

At Longevity Summer Camp 2022 (a program of the Longevity Biotech Fellowship), we held a discussion to identify systemic deficiencies in the longevity field (termed longevity bottlenecks). We discovered that while it was clear that many bottlenecks existed, there was no obvious way to assess their severity and relative importance. We also found that no established methodology to rigorously identify and prioritize bottlenecks existed. We therefore set out to 1) establish a systematic way to identify bottlenecks of a field and 2) perform the analysis on the longevity field.

Instead of using a think tank approach where a small group of people gather, analyze existing data, and produce projections and recommendations, we opted to directly identify bottlenecks by surveying a broad set of participants across different sectors of the longevity field. To avoid theorizing and instead obtain experience-rooted answers, we asked for bottlenecks that participants experience **in their own work**, with the goal of identifying common trends in people’s bottlenecks in a representative and unbiased way. We also asked for participant’s most needed solutions, offering insight into each respondents’ circumstances to provide higher specificity. Lastly, we asked participants to evaluate the perceived efficacy of interventions that they had professional expertise in, and to judge these interventions by the relative severity of bottlenecks affecting them.

## Results

### Participants

We contacted approximately 1000 participants in academia, biotech, pharma, policy, media and other areas of the longevity field between December 2022 and February 2023 and asked them to fill out an online survey (full survey in **Fig. S1**). Of those contacted, 395 completed the survey (see Supplementary File 1 for the full survey). No personally identifiable information was collected and as such, the analysis was blinded. Among the participants, the most frequent role was Academic research scientist (Postdocs or PhD/MSc/BSc student), followed by Principal Investigators/Professors, Entrepreneurs, Biotech Research Scientists and other occupations **(Fig. 1A)**. To reflect the interdisciplinary nature of the field, we allowed each participant to select more than one role, and as such the total number of responses in Fig. 1A exceeds the number of participants. Nevertheless, most participants (74% of total) selected a single role. The participant pool was well diversified in tenure in the field, containing roughly equal numbers of newcomers as well as veterans **(Fig. 1B)**. Finally, the most common personal longevity goal was to live until 100 years of age **(Fig. 1C)**, while almost as many people wish to live indefinitely or as long as they could remain healthy. Some wish to live until a certain event occurs, such as loss of their family or loss of their job.

**Figure 1.**
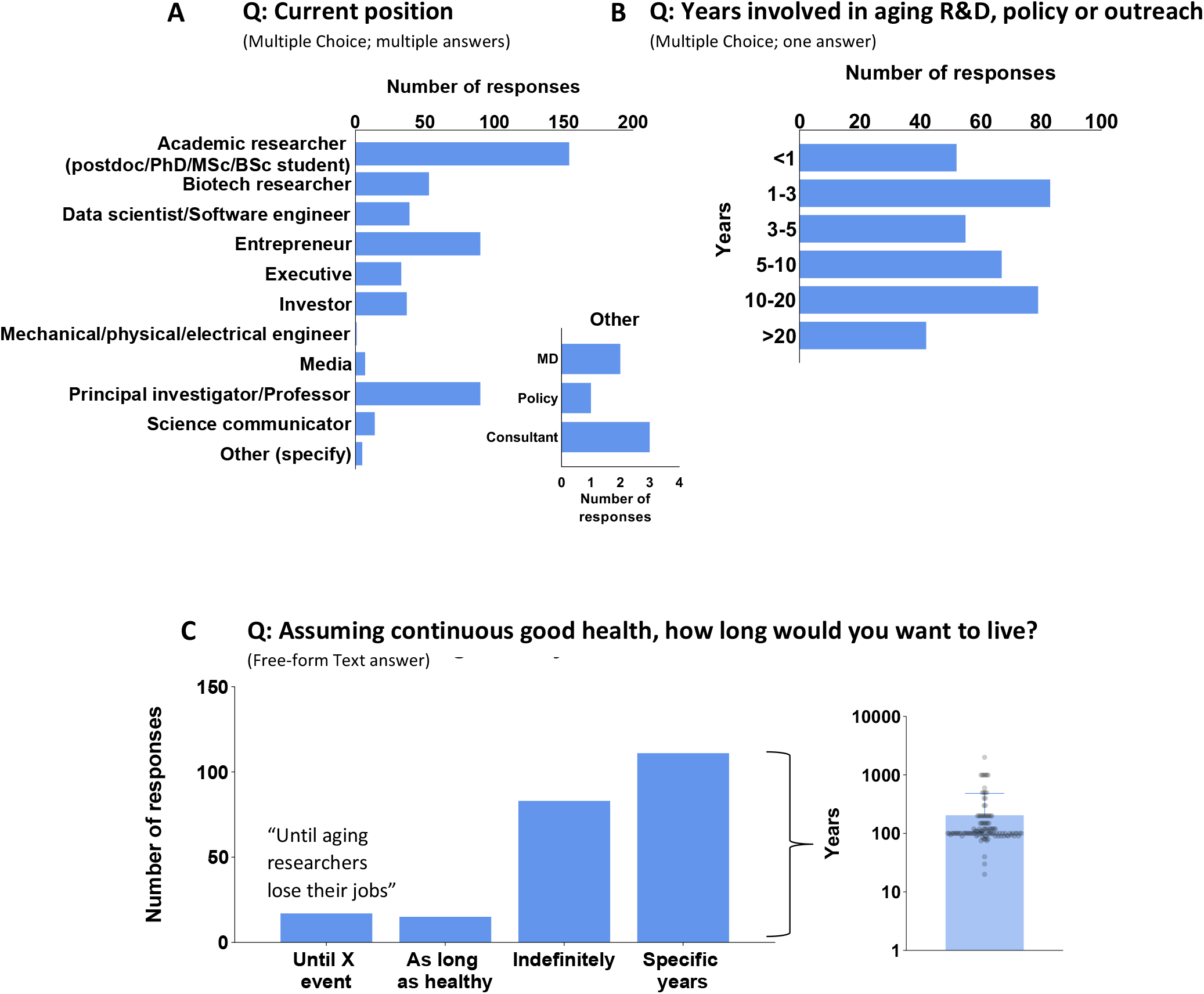
Background and personal longevity goals or participants. Question listed above each graph. **(A)** Current position of participants (Multiple choice question, multiple selections allowed). Inset shows categories listed in “Other”. **(B)** Number of years participants have been involved in the longevity industry (single choice question). **(C)** Personal longevity goal of participants (free-form question, with one example provided). See Figure S1 for the full survey.

### Baseline attitude towards the current state of the longevity industry

To estimate the perceived current rate of progress, we asked, “*At current rates of progress, how many years of life do you think we will add to the average lifespan of generally healthy people in 5, 10, 25 years, assuming existing bottlenecks largely stay the same?*” The average of estimates to the questions were 2.1 years added in 5 years of research; 5.8 years added in 10 years of research; and 18.2 years added in 25 years of research **(Fig. 2A)**. Therefore, most participants do not think we are close to achieving “longevity escape velocity” – the point where one year of research will produce an average increase of 1 year or more in healthy lifespan. Furthermore, here as well as throughout the study, there is substantial disagreement in the answers, as variation between individual estimates was large, varying up to three orders of magnitude.

**Figure 2.**
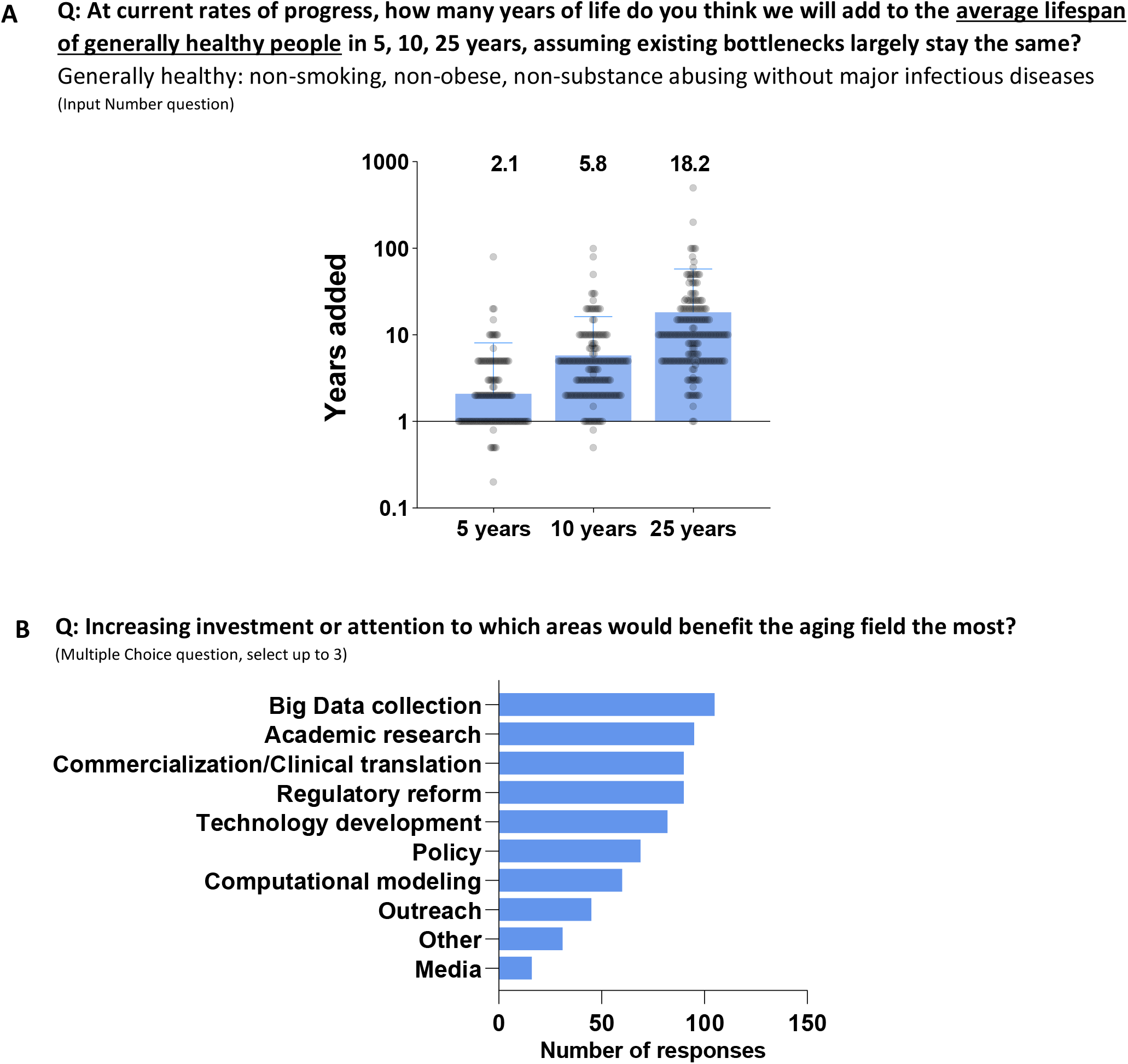
Perception about the current state of the field. Questions are listed above each graph. **(A)** Perceived rate of progress given current state of the field, displayed at log10 scale. Some values lie between 0.1 and 1 because some participants answered in negative numbers. **(B)** Perception regarding where additional investment would most benefit the field (multiple choice question, up to three answers). See Table S1 for answers listed as “Other”.

To estimate the perception about the area where investment would result in the biggest benefit to the aging field, we asked “*Increasing investment or attention to which areas would benefit the aging field the most*?” Surprisingly, Big Data collection was rated as the most important area to invest in, receiving over 100 votes, while Media was considered the least important, receiving 10-fold fewer votes **(Fig. 2B)**.

### Bottlenecks

To estimate the bottlenecks to progress in the longevity field, we asked “*In your work in aging, what are the biggest bottlenecks you currently face? Be as specific as you can*.” with up to three free-form text answer boxes. We received close to 1000 answers. To analyze this dataset, we established a two-tiered system, where we first identified and defined individual Bottlenecks, then grouped them in Categories, and finally tagged every answer with one or more Bottlenecks **(see Fig. 3A; *Methods*)**. For clarity, we denote these Bottlenecks by their ID, enclosed in brackets (e.g. [Bottleneck]).

**Figure 3.**
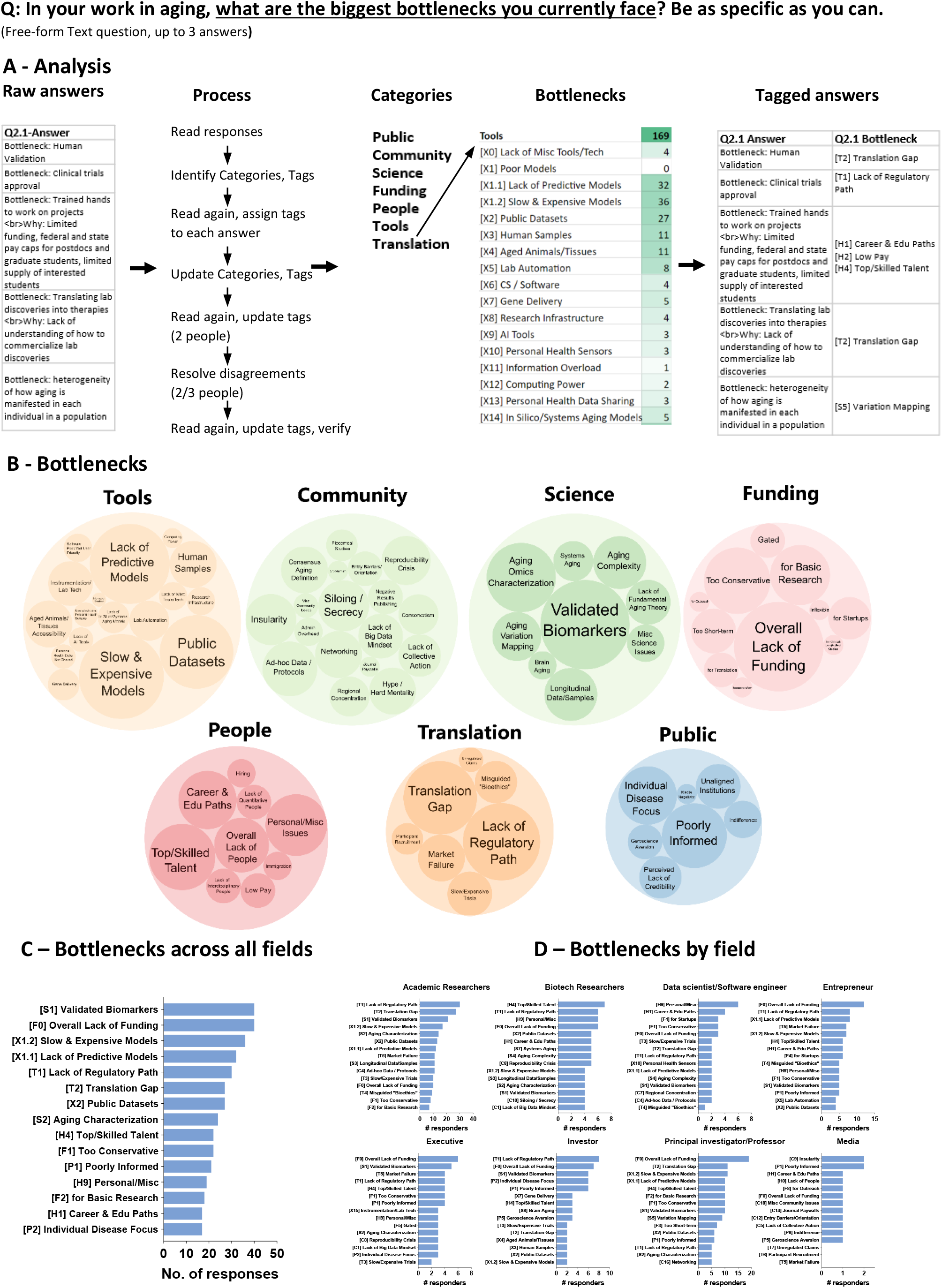
Bottlenecks of participants. Short-form question listed above panel A (free-form text, up to 3 answers). **(A)** Analysis of the free-form text data to generate Categories and Bottlenecks tags. **(B)** Summary of bottlenecks listed by participants. For both Categories and Bottlenecks, the area of the bubble is proportional to the number of answers. **(C)** Top 15 bottlenecks across all answers. **(D)** Top 15 profession-specific bottlenecks. The [XX] tag in front of each Bottleneck description represents its unique qualifier in our bottlenecks classification system. See Table S2 for full list of all Bottlenecks.

The most frequently identified bottlenecks across all participants were lack of [Validated Biomarkers] and [Overall Lack of Funding], receiving 40 mentions each. This was followed by [Slow & Expensive Models], a [Lack of Predictive Models] (36 and 32 mentions) and a [Lack of Regulatory Path] (30 mentions) **(Fig. 3B-C)**. However, the major bottlenecks varied substantially by profession **(Fig. 3D)**. [Overall Lack of Funding] is the biggest bottleneck for Entrepreneurs, Executives and Principal Investigators/Professors, whereas [Lack of Regulatory Path] is the major issue for Academic Researchers and Investors. Finally, for Biotech researchers, the major bottleneck is the lack of [Top/Skilled Talent], for Data Scientists and Software Engineers, it is [Personal/Misc.] circumstances and lack of [Career & Edu Paths], and for Media it is [Insularity of the community] and [Poorly Informed] public.

### Solutions

To find out what solutions are most needed, we asked “*If it was available, which one tool, resource, or regulatory/social/other type of change would have the biggest positive impact on your work*?” and provided one free-form text box for answers. We repeated the analysis process described in Fig. 3A to establish a categorized two-tiered system for Solutions. Interestingly, despite being a free-form answer in a different context, the most mentioned solution across all participants was increased availability of [Public Datasets] **(Fig. 4A-B)**. This matches the result of our question *“Increasing investment or attention to which areas would benefit the aging field the most?”* (Fig. 2B). The next most wanted solutions were the establishment of defined [Regulatory Path], [Overall More Funding], establishment of [Validated Biomarkers], and improved [Networking] opportunities **(Fig. 4B)**. As before, most wanted solutions differed by a participant’s field. [Public Datasets] was named as the top solution for Academic Researchers and featured in the top 10 for most other professions. [Overall More Funding] was the most sought solution for Data Scientists/Software Engineers and Principal Investigators/Professors, while more defined [Regulatory Path] was the most needed solution for Biotech Researchers, Entrepreneurs, Executives and Investors. Finally, for people in Media, more [Public Outreach] and Communications Training] for scientists were the most wanted solutions **(Fig. 4C)**.

**Figure 4.**
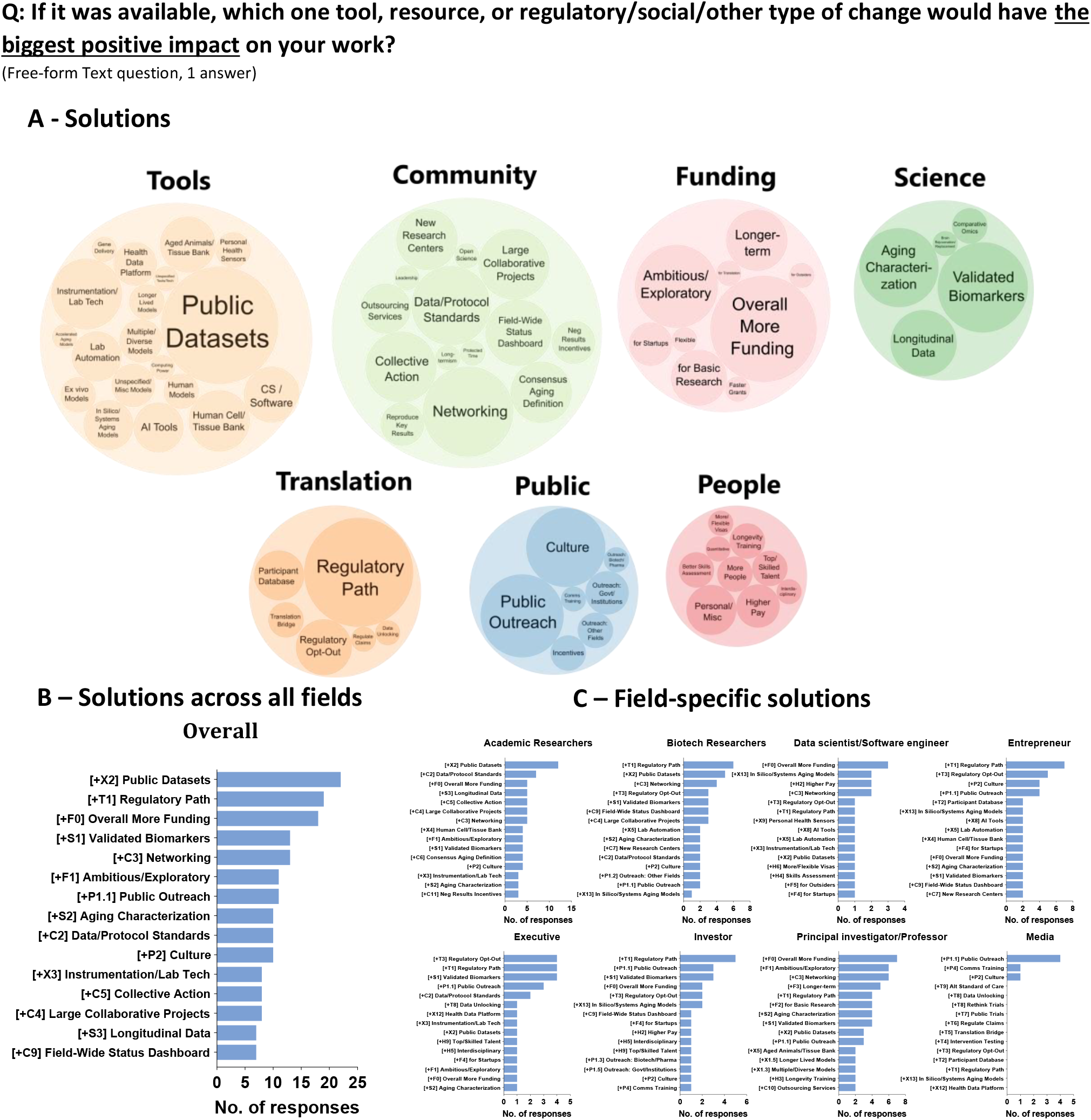
Solutions most needed by participants. Short-form question listed above panel A (free-form text, 1 answer). **(A)** Summary of solutions listed by participants. For both Categories and Solutions, the area of the bubble is proportional to the number of answers. **(B)** Top 15 solutions across all answers. **(C)** Top 15 profession-specific solutions. The [XX] tag in front of each Solutions description represents its unique qualifier in our classification system. See Table S3 for full list of all Solutions.

### Perceived efficacy of interventions and their associated bottlenecks

In our effort to estimate field-wide bottlenecks, in addition to the free-form text analysis above, we designed a top-down approach centered around individual well known interventions and their associated bottlenecks. In this approach, we asked participants to rate a predefined list of interventions for the number of years they think the intervention will add to healthy life after 5 years, 10 years and 25 years of continued research. This approach allowed us to compare the widely distinct interventions within a single dimension in the short, medium and long-term. To prevent theorizing, we first asked which of the listed interventions participants were familiar with and directed them to evaluate only these interventions (self-defined expert group) **(Fig. 5A**,**B and Fig. S2A, Fig. S3)**. We also allowed participants to separately evaluate the interventions they were not familiar with after they completed the previous step (self-defined non-expert group), and analyzed data from the two groups separately. In the short term (5 years), rapamycin supplementation and calorie restriction are considered as the most efficacious interventions, both thought to add about 2.5 years of healthy life **(Fig. 5B)**. In the medium term (10 years), a number of other interventions, including telomere extension, reprogramming, stem-cell based and genetic medicine-based therapies (gene therapies and gene editing) are considered more effective, with each thought to add about 5 years to healthy life. In the long term (25 years), reprogramming, genetic medicine and organ replacement are considered most promising **(Fig. 5B)**. Interestingly, sirtuin targeting therapies and NAD targeting therapies are perceived to be poorly effective in every timeframe **(Fig. 5B)**. These results were mostly replicated between both self-defined expert and non-expert groups **(Fig. S2B)**.

**Figure 5.**
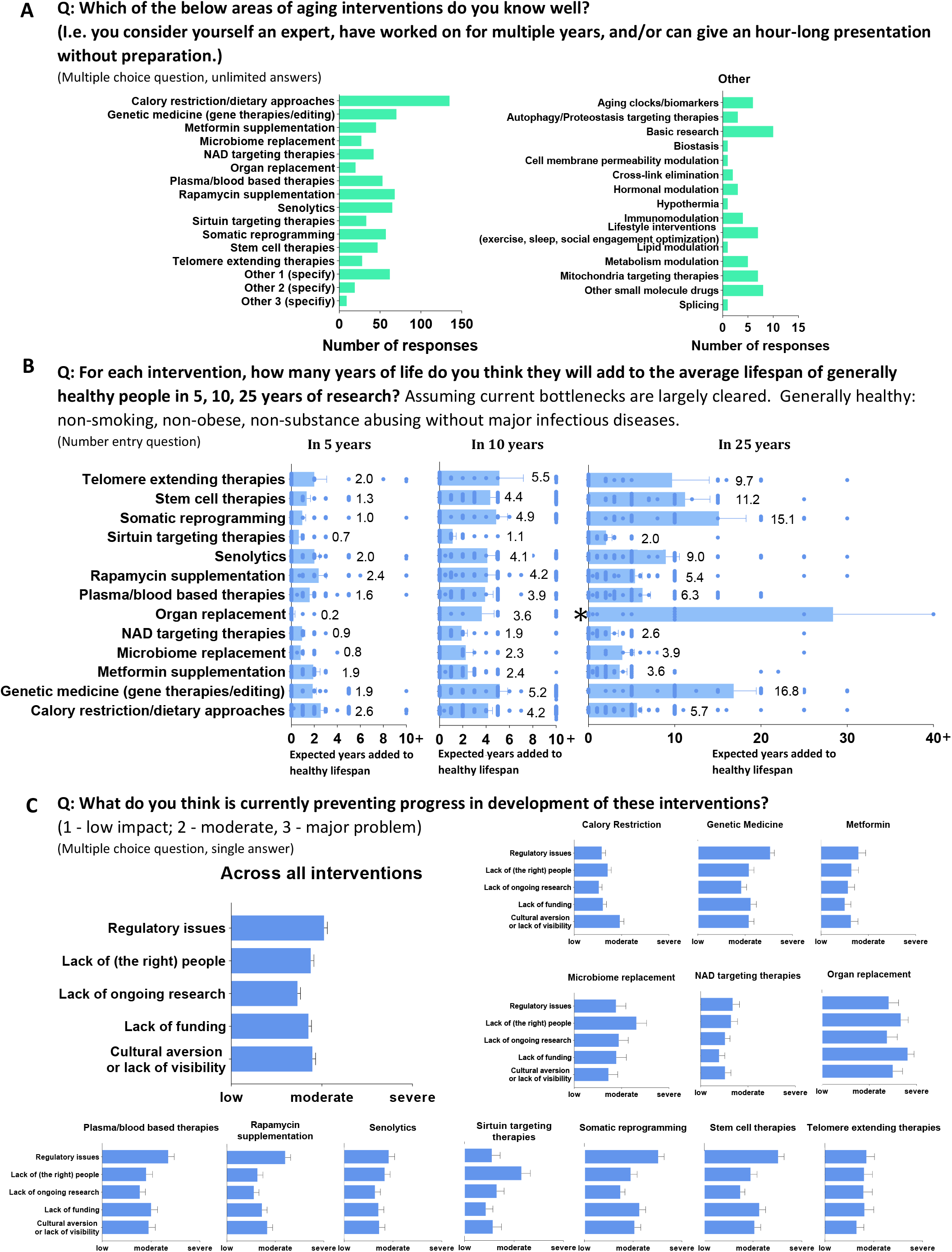
Bottlenecks by interventions. The question (in short form) listed above each panel. **(A)** Interventions that participants self-identified as being expert at and subsequently evaluated. In panels B-C, participants evaluated only the interventions they individually selected in Panel A. Inset – interventions listed under “Other”. **(B)** Perceived efficacy of interventions across three timescales. Each dot represents one individual estimate. Data plotted as linear graphs with X axis range centered around the mean values to allow comparison of means of different interventions. Dots on the very right edge of the graph represent estimates of 10 years or more (left and middle) or 40 years or more (right). *Note that the mean value for organ replacement on the right graph is substantially influenced by a single estimate of 200 years; removal of that estimate places the expected years of life added at an average of 16.1 years (see Fig. S2). **(C)** Perceived severity of impact of five categories of bottlenecks on interventions. (B-C) Error bars represent standard error of the mean (SEM). For interventions evaluated by self-identified non-experts, see Figure S2. For evaluations of interventions listed as Other, see Figure S3.

Finally, we asked how much certain bottlenecks affect progress in each intervention. To keep the questionnaire at a manageable length, we offered 5 categories of bottlenecks as possible answers: Regulatory Issues, Lack of (the right) People, Lack of Ongoing Research, Lack of Funding, and Cultural Aversion/Lack of Visibility. Participants were asked to rate these categories on a 3-point scale, ranging from low, moderate and severe impact for the intervention in question. Across all interventions, the category considered most severe was Regulatory Issues **(Fig. 5C)**. Regulatory Issues were estimated as the most severe bottleneck for genetic medicine, rapamycin and metformin supplementation, reprogramming, senolytics, NAD targeting therapies, stem cell therapies, and telomere extending therapies. For other interventions, the perceived most severe bottlenecks differed. Interestingly, all factors were estimated as close to severe for organ replacement, and Lack of (the right) People was estimated as the biggest issue for sirtuin targeting therapies and Microbiome replacement therapies **(Fig. 5C, S2)**.

## Discussion

### Lack of validated biomarkers and funding

The two biggest bottlenecks across all respondents were a lack of [Validated Biomarkers] and an [Overall Lack of Funding] (40 votes each). Two conclusions can be drawn from this. First, while substantial new funding has entered the field over the past five years, it does not seem to have reached most participants, possibly because these sources of funding are concentrated in a few large institutions or corporations. Second, while much progress has been made in the development of aging biomarkers, many participants do not feel that these biomarkers are sufficiently validated to be used as predictive endpoints in clinical trials. Therefore, to accelerate progress across the field, it may be more effective for investigators working on biomarkers to focus on the clinical validation of existing biomarkers rather than the development of new biomarkers.

### Poor models

The third and fourth most commonly identified bottlenecks were [Slow and Expensive Models] and the [Lack of Predictive Models]. Specifically, this means that the animal, cell and/or tissue culture models in use are perceived to be too slow, too expensive and/or not sufficiently predictive of human outcomes. If these two bottlenecks were combined into one (a general lack of good models), this would be by far the most commonly named bottleneck. These answers indicate that as a field, we need to improve the economics of the existing models to lower their cost, establish ways to identify results from them faster, and develop rigorous methodology to identify where models are predictive of human biology and where they are not. The fact that all drugs are currently tested in models of some kind as a precondition to human trials, with very little systematic confidence in the predictive power of those models, should be a major stain to the image of biologists everywhere, but is particularly dangerous to longevity researchers due to the longer timescales associated with studies.

### Regulatory deficiencies

The lack of regulatory paths was listed as a major bottleneck by multiple professions. Many answers also noted that aging not being classified as a disease is a substantial contributor to this problem. These answers should be placed in context. Classification of aging as a disease may indeed improve clarity in regulation, but multiple endpoints that are commonly employed in animal longevity studies (lifespan, all-cause mortality, and aspects of the frailty index) are in fact recognized by the FDA (3). However, these metrics are not commonly employed in human geroscience trials because of the cost and timescales required to observe statistically significant changes. For this reason, use of biomarkers predictive of longevity instead of these direct metrics would be advantageous, but as discussed previously, many survey participants do not see current biomarkers as sufficiently validated for translation. Given this lack of validation, it is not surprising that biomarkers are not currently accepted as endpoints by the FDA. Identifying robust, validated biomarkers that track or predict these three FDA-recognized metrics relevant to longevity may be a more effective path to regulatory clarity than achieving classification of aging as a disease.

### A lack of established paths for newcomers

While [Overall Lack of Funding], [Lack of Regulatory Path], and lack of [Validated Biomarkers] were listed as top bottlenecks for most professions, for Data Scientists and Software Engineers, the main bottlenecks were [Personal/Miscellaneous] factors, along with lack of [Career & Educational Paths]. For us at the Longevity Biotech Fellowship, this is a well-recognized pattern. Migration of people from other fields into longevity has ramped up in the past 5 years, including from software engineering, data science, and other fields primarily involved with data and coding. However, people from these fields commonly face difficulties in entering longevity due to a lack of accessible and rigorous longevity educational resources, paucity of formal degrees in longevity, immigration barriers, and lower salaries compared to their industries of origin. While increasing pay may be challenging given that [Overall Lack of Funding] was tied as the #1 bottleneck, other entry barriers can be reasonably addressed with community efforts. We encourage readers to join us in these efforts (see *Implementing solutions*).

### Efficacy of interventions

While we asked participants to estimate the number of years added to healthy lifespan for different interventions, the goal of this question was not to attempt to forecast the efficacy of each intervention. In fact, the absolute values of these predictions may make for good entertainment by the time we reach the listed timepoints (5, 10 and 25 years from 2023). Rather, our goal was to identify the *current sentiment* of well-informed participants of the longevity field to these interventions, reasoning that these estimates reflect the *relative* expected future efficacy of different interventions based on currently available animal and human data. We identified rapamycin supplementation and calorie restriction as the interventions considered most efficacious in the near term and reprogramming, genetic medicine and organ replacement as those considered most efficacious in the long-term. Additionally, we found that there is substantial skepticism towards the efficacy of NAD and sirtuin targeting therapies. For the latter, different possible interpretations should be considered. There is substantial literature indicating the efficacy of NAD and sirtuin targeting approaches in increasing health (*4, 5*), although some of the interpretations from the studies on sirtuins are contested (*6*). While animal evidence exists for lifespan increase for both NAD and sirtuin targeted interventions, the effect sizes have been moderate or sex-specific (*7, 8*) (until the recent work published by Roichman *et al*. (*9*)). Therefore, one interpretation is that participants may have focused on the lifespan data in their answers. Another interpretation is that NAD and sirtuin targeting therapies are thought to generally be of low efficacy, reflecting the historical controversy around these interventions (*10*).

### Solutions Throughline: a need for a data-first science

The most voted solution across two separate questions (Fig. 2B and 4B) was the development and sharing of large-scale public datasets. Many answers that mentioned this also listed other bottlenecks that prevent data sharing or unified data analysis as major issues. Summarizing these and other answers, one throughline emerges from this survey: we need a fundamental, data-oriented evolution in the way we do science. This was well captured in one answer: “*Bottleneck: Artisanal science institutions. Why: R&D is mostly still done following century old methods of manual labor and hypothesis and publications ridden with narrative fallacy. Our institutions and funding are setup to maintain it this way. We need to switch to automation, massive empirical data collection, and computational modeling*.*”* In various forms, this sentiment was echoed by many participants, who added the bottlenecks of [Reproducibility Crisis] and lack of [Negative Results Publishing], data [Siloing/Secrecy], [Insularity], [Ad-hoc Data/Protocols] and [Lack of Big Data Mindset] to the above.

### Implementing solutions

The real issue with most of the identified bottlenecks is not the lack of possible solutions, but the current lack of directed efforts to achieve them in any reasonable timeframe. Systematic issues in our tools, approaches and conventions in performing and disseminating research receive a fraction of the attention and investment that is spent on the development of individual new drugs. Yet, major breakthroughs in biology have mostly come from the development of new technologies that have often required decades-long concerted efforts (for example, next-generation sequencing and gene editing). Some of the bottlenecks identified above are well known and difficult to solve technically, societally or financially. Others, however, are tractable and have possible solutions that could be immediately implemented. Examples of such solutions from our data include:

- Data/Protocol Standards Consortium
- Accessible longevity education and training programs
- A central investor-scientist-entrepreneur networking hub
- A field-wide progress dashboard (from research hypothesis through to clinical trial outcomes)
- Incentives or mechanisms for publishing negative results (i.e. through a web portal, journal, etc.)
- Automated AI tools for publication quality control
- Aged animal and human tissue biobanks
- Machine-readable datasets/papers
- Aging Definitions Consortium: formalizing terms and definitions used in longevity for improved communication and codifying them with National Institute of Standards and Technology
- Incentives for large-scale adoption of Open Access and preprint servers for removal of paywalls
- “Meta” database of databases for easy automated discovery
- “Essential Tools for Longevity Scientists” articles, sites and resources.
- Shifting grant allocation to more large-scale public data collection projects (akin to the Human Genome Project or large physics collaborative projects).
- Stimulating competition through more longevity prizes (akin to XPRIZE)
- Incentivize technology development over discovery-oriented research (i.e. RFAs for tech and dataset development)
- Establish a recognized Longevity R&D Advising group to guide investors and funders to high-impact, underserved areas of research and technology development.
- Etc.

These solutions are tractable with current technologies and levels of funding, but need community organization and buy-in. In order to drive development of these and other solutions, we at the Longevity Biotech Fellowship are organizing working groups (including both for-profit and nonprofit ventures) to tackle individual bottlenecks. If you wish to join or nucleate a group, please contact the corresponding authors.

## Methods

### Survey methods

The full survey is listed as Fig. S1 and full study data is listed as Supplementary File 1, as well as available in a browsable form at https://pmurs.github.io/bottlenecks/#/. The survey was administered using Qualtrix. The study plan was submitted for evaluation by the Harvard University-Area Committee on the Use of Human Subjects and exempted from IRB review based on regulations found at 45 CFR 46.104(d) (2) (Exemption IRB22-1588). The study target population was adult (18+ year old) individuals working in or associated with the longevity field, and formal consent was asked at the beginning of the survey to restrict the participant pool to this population. No personally identifiable data was collected. For multiple choice questions, the order of choice answers was randomized among different participants to prevent bias.

### Collection

Study data was collected between December 2022 and February 2023. To ensure quality of the data, most participants were invited to the survey through a unique randomized study link that expired after completion of the survey. Occasionally, participants opted to recruit further participants and received additional unique randomized links from the study PI. A small minority of answers were received through a general survey link shared to small groups of people. Survey participants received two follow-up requests for study completion. Participants were contacted directly by the study PI through email, LinkedIn or text and drawn from a semi-random pool of academics, academic research scientists, biotech or industry research scientists, entrepreneurs, investors and other sectors of people known to be associated with the longevity industry by the study PI. Approximately 1000 people were contacted, 800 people opened the survey and approximately 400 people completed the survey. After removal of duplicates, a dataset of 395 unique sets of answers were used for analysis. No link between unique randomized study link and participant IDs or emails were retained after data collection and quality control were completed.

### Analysis

For Figures 1C, 3, 4 and 5A, answers from free-form text questions were analyzed by the study authors and each answer assigned to categories listed in corresponding figures (see Fig. 3A). Each answer was analyzed and categorized by a minimum of two evaluators and reviewed by a third evaluator in the case of disagreement. For Figures 3 and 4, a two-tiered Bottlenecks and Solutions categorization system was first constructed consisting of 1. Categories and 2. Bottlenecks/Solutions, whereby each Bottleneck or Solution falls under a Category. Each answer was then tagged for one or more Bottlenecks/Solutions respectively. This categorization was established through the following method. First, answers from free-form text were read by 3 separate evaluators and approximate Categories and Bottlenecks/Solutions were independently proposed by each evaluator for each response. The common overlap between proposals was conserved, and disagreements were resolved via discussion, establishing a preliminary classification of Categories and Bottlenecks/Solutions. The answers were read again by three evaluators and each answer tagged with one or more Bottleneck or Solution. Then, the classification of two evaluators was compared, and the Categories and Bottlenecks/Solutions tags were further refined. Then, free-form answers were read again by two evaluators, updated Categories and Bottlenecks/Solutions tags were applied to each answer, the results of two evaluators were compared and the disagreements were resolved by the third evaluators assessment. Finally, answers were read again by a single evaluator, all updated Category and Bottleneck/Solutions tags were applied, and answers were verified for the correctness of the tags. Finally, the data were analyzed in Microsoft Excel and visualized using GraphPad Prism.

### Possible Limitations

This study had some limitations. First, we are not aware of any field-wide data sets listing the number of people involved in different sectors of the longevity field - therefore it is difficult to estimate whether our participant group was a representative sample of the longevity industry. In our dataset, academic researchers, professors and principal investigators and entrepreneurs were the most populous respondent groups. As such, all analysis sets that pooled data from the whole dataset were majorly influenced by these groups. For this reason, we provide sub-group specific analysis where particularly relevant. Secondly, most participants were sourced from the United States; therefore, the results above may be specific to this country. Furthermore, we did not investigate geographic or age-specific differences in answers because we chose not to collect personally identifiable data, including geographic location or age of the participants. Finally, we chose to allow participants to self-identify across multiple groups rather than forcing a single identification to accurately reflect the complexity of people’s roles in this field. As such, the participants who self-identified across multiple groups contributed to the analysis of answers from all groups that they identified as a part of. Nevertheless, most participants (74%) self-identified within one group only and substantial differences in top bottlenecks and wanted solutions were present between different groups, indicating that the above did not have an overt effect on our analysis.

### Followup Studies

During the process of analyzing the survey data we recognized numerous improvements that could be made to future iterations of the survey, for instance: 1) differentiating between intervention targets, specific therapies and intervention modalities, 2) A pre-survey questionnaire to establish a more comprehensive list of intervention targets/therapies and 3) increasing outreach to improve sample size, relative representation of respondent types, and geographical diversity. We plan to repeat this survey at regular intervals to continue to guide resource allocation in the field. The authors welcome feedback and ideas for improvements to future surveys.

## Supporting information

Supplementary File 1 - Survey data and analysis

## Acknowledgements

We would like to thank Todd Huffman for hosting the Longevity Summer Camp 2022 conference and Prof. Amy Wagers, Eva Tomczyk and Kristen Elwell from Harvard University for help with navigation of the IRB review process.

## Authors Contributions

Mark Hamalainen (MH) organized Longevity Summer Camp in 2022; MH and Michael Florea (MF) organized the original bottlenecks discussion group; all authors participated in the discussion and development of the study format and questions; MF, MH and Patrick Seebold (PS) coded the survey; MF and PS organized IRB review; MF, MH and Nathan Cheng (NC) collected answers; MF, MH and PS analyzed the data; MF wrote the paper; MF, MH and PS edited the paper; MF, Paul Murray (PM) and NC created the figures; PM created the online data portal.

## Supplementary Information

**Figure S1.**
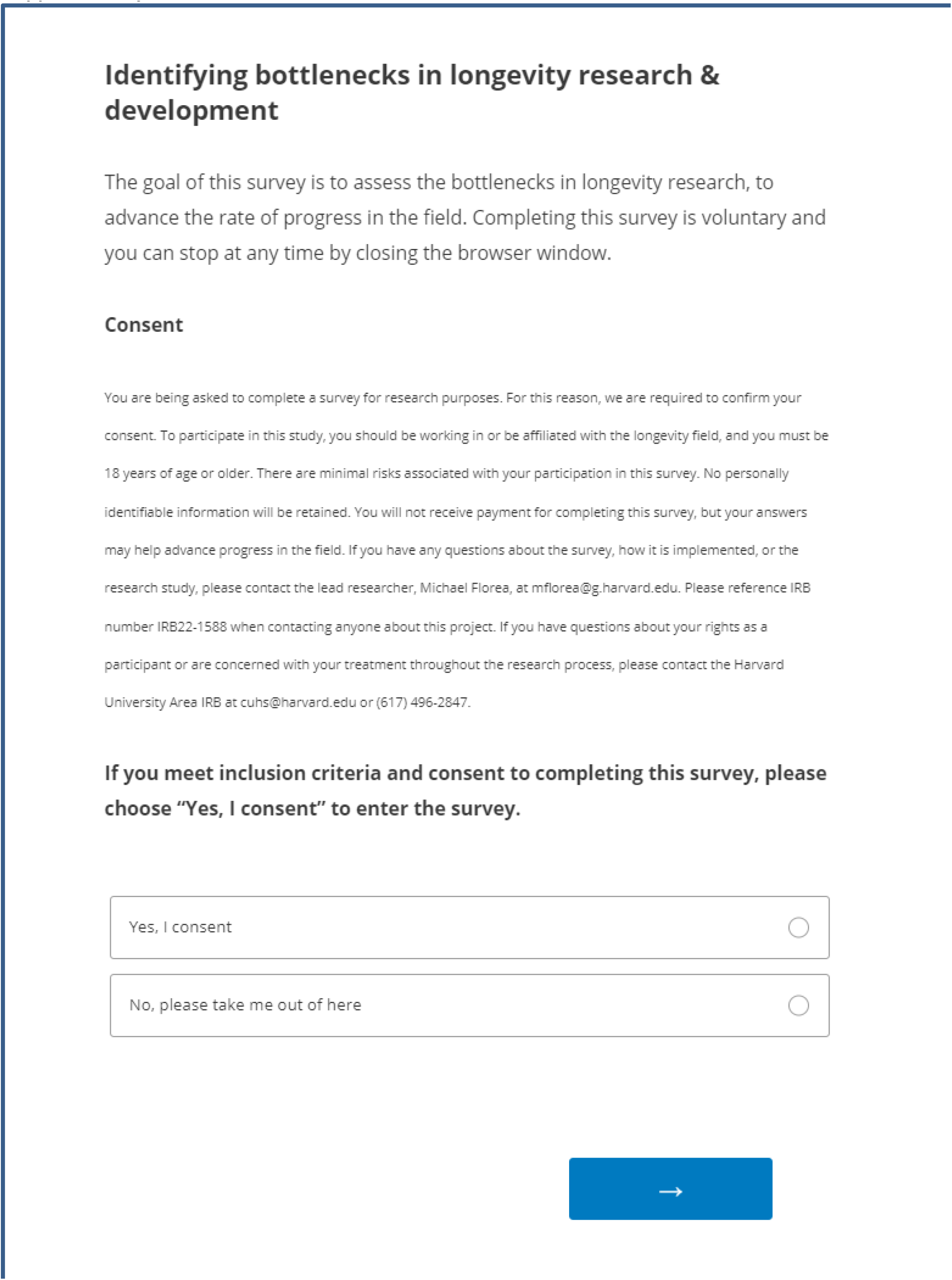
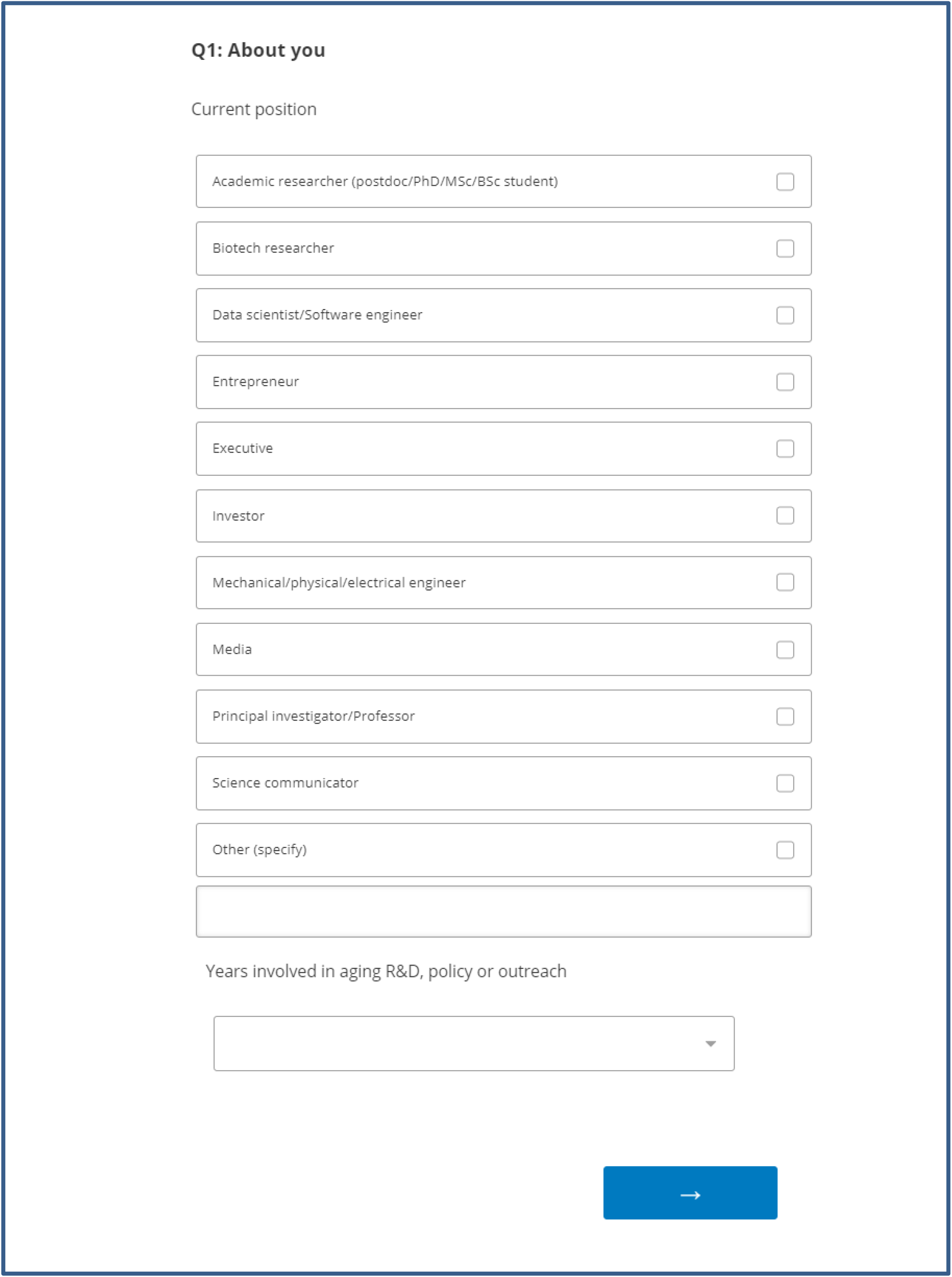
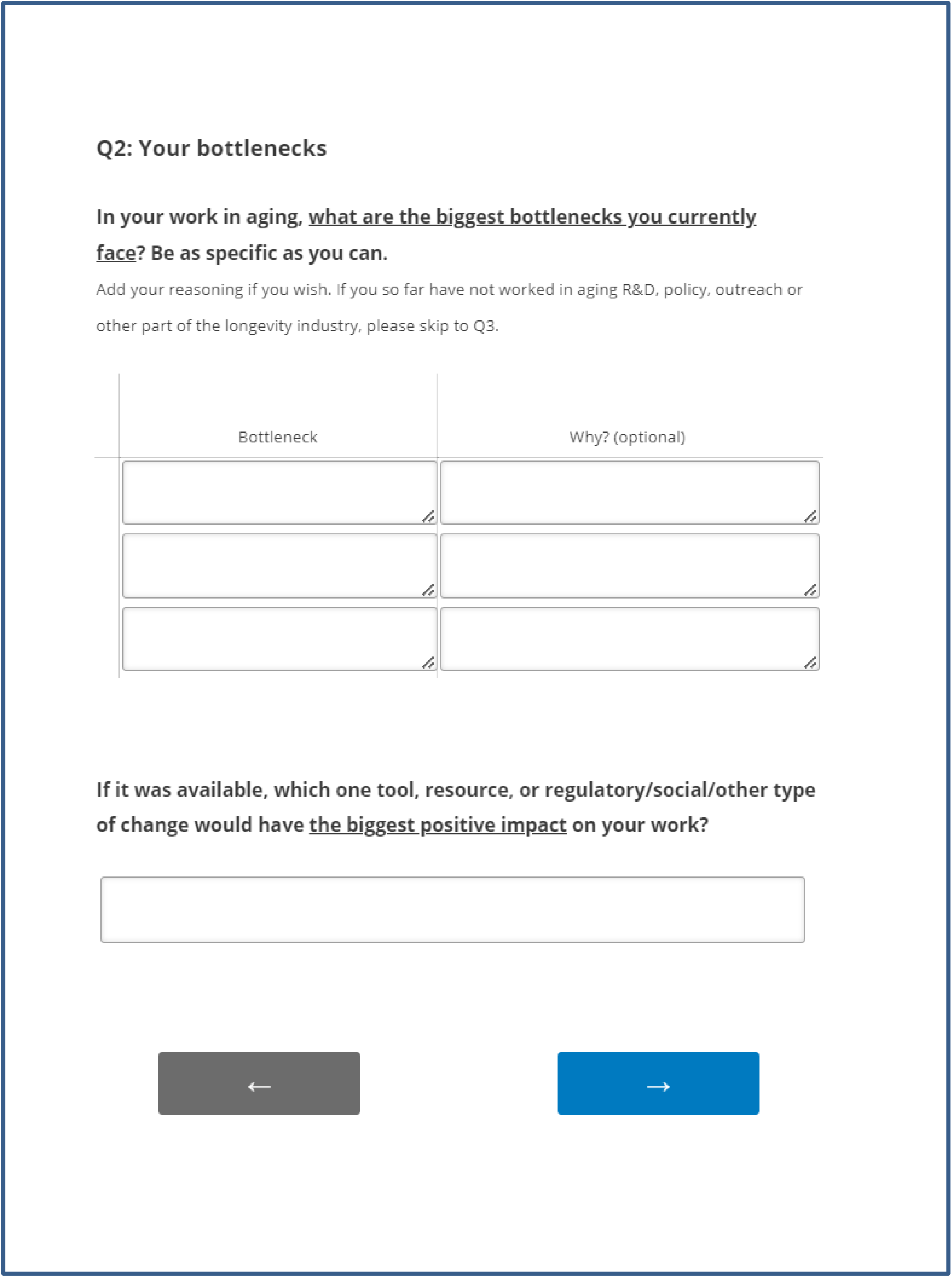
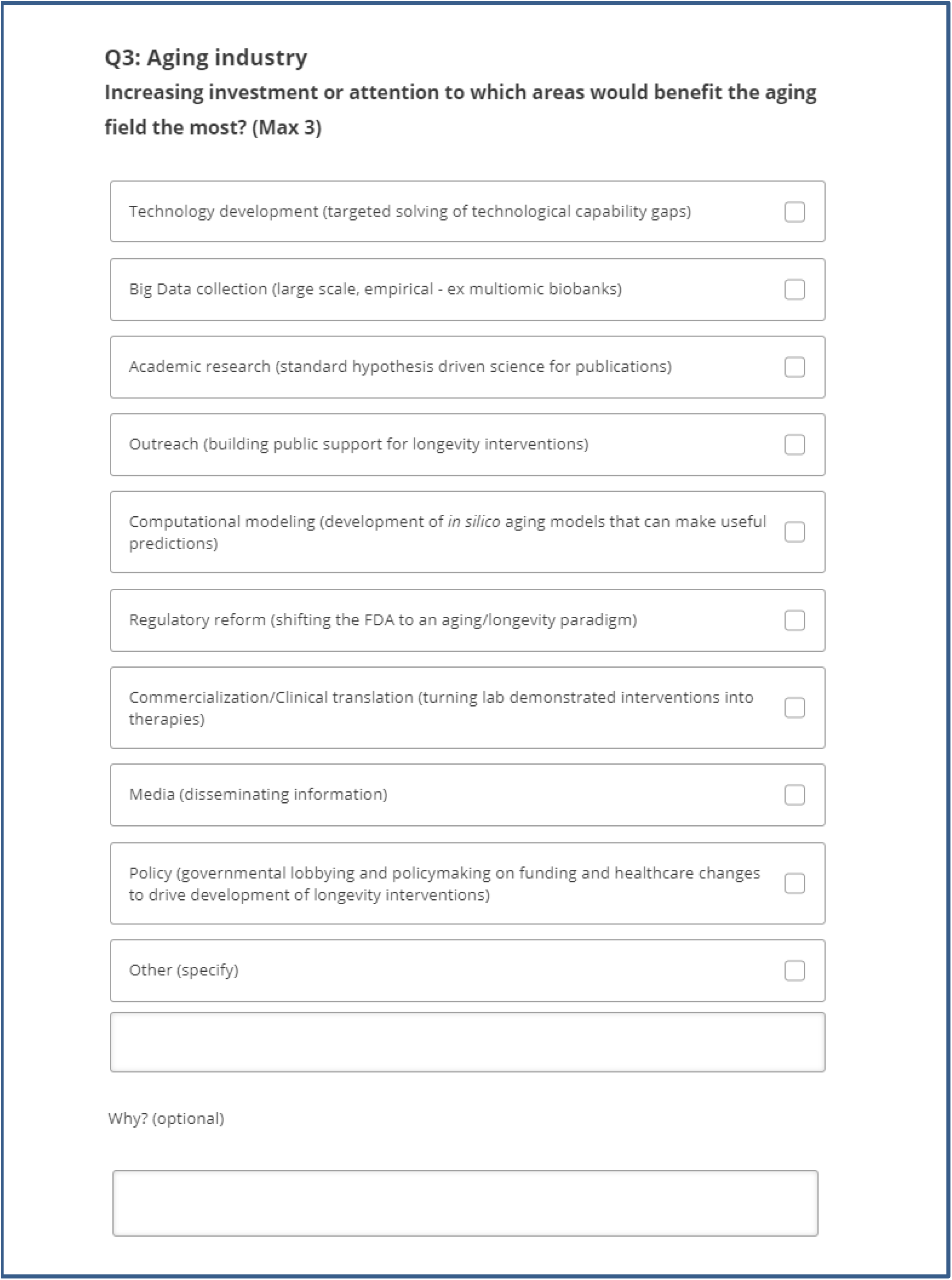
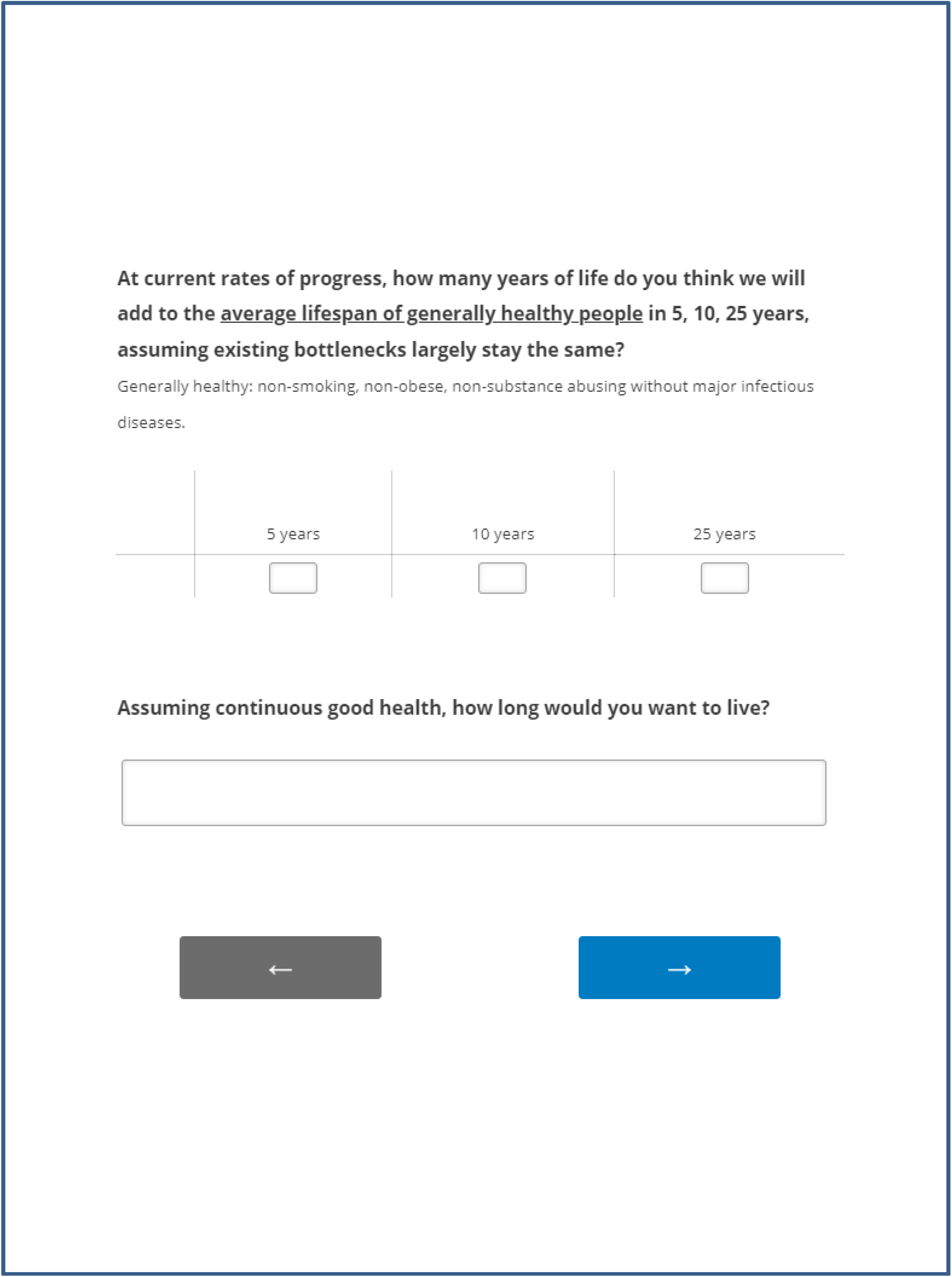
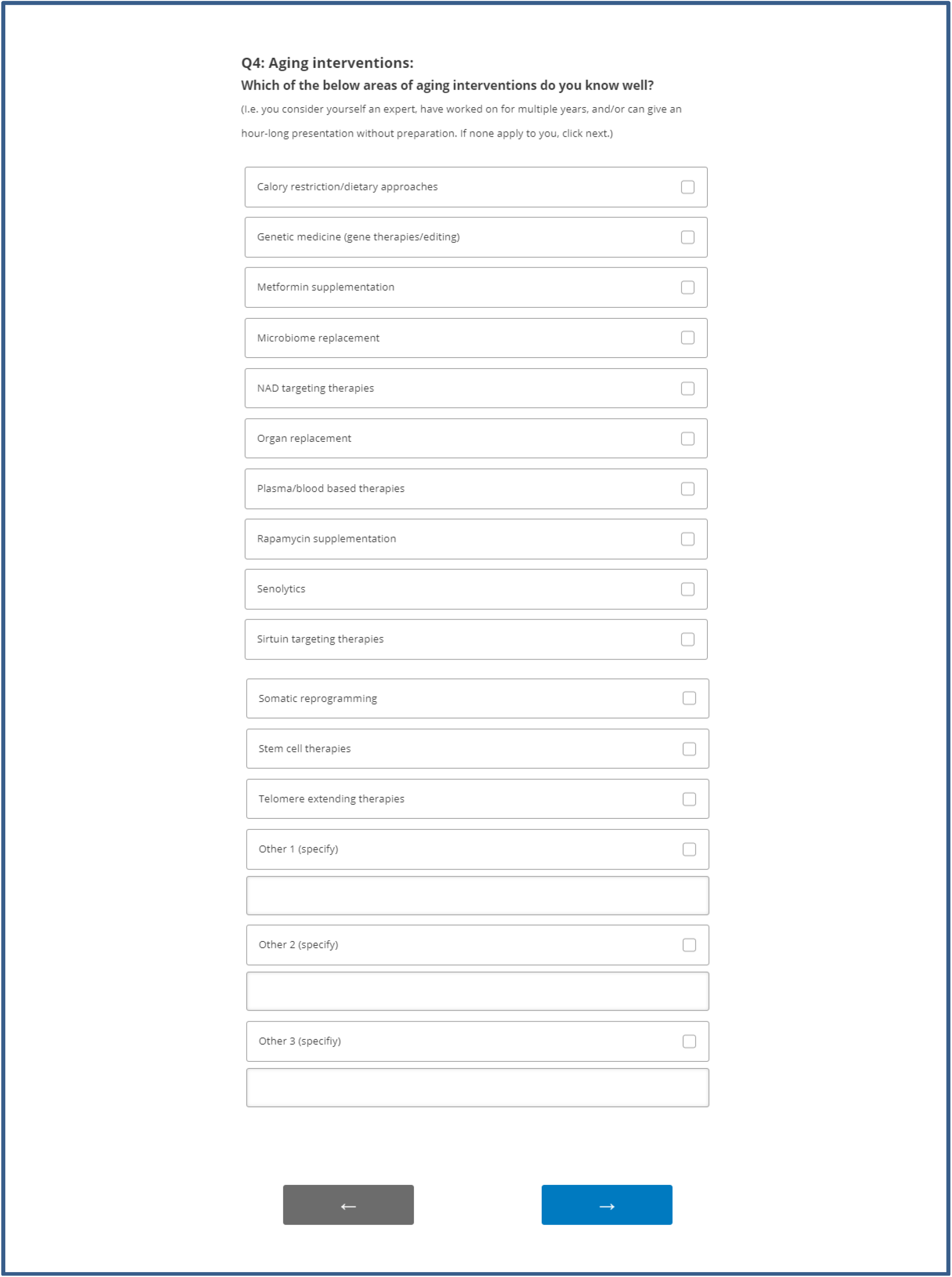
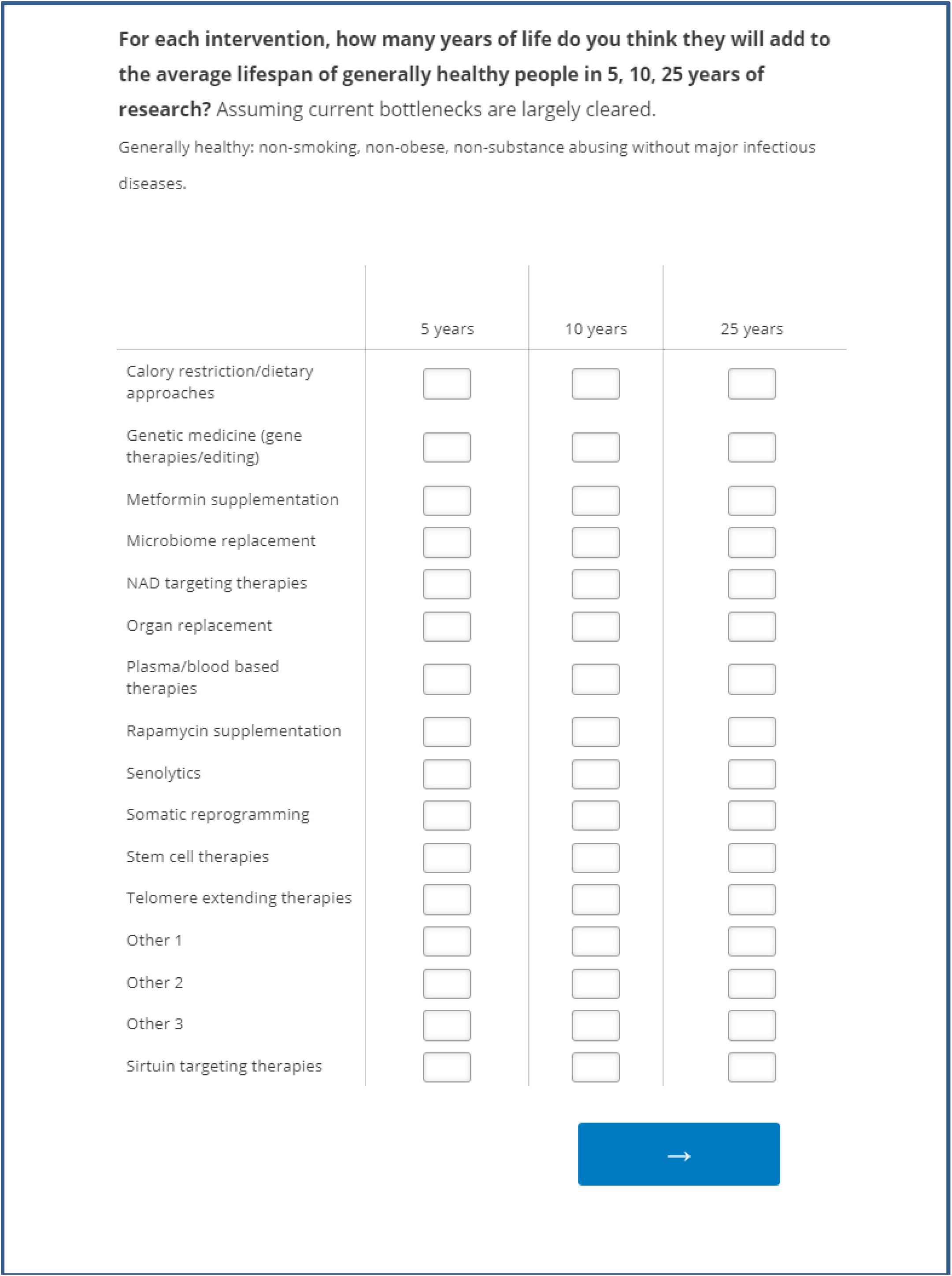
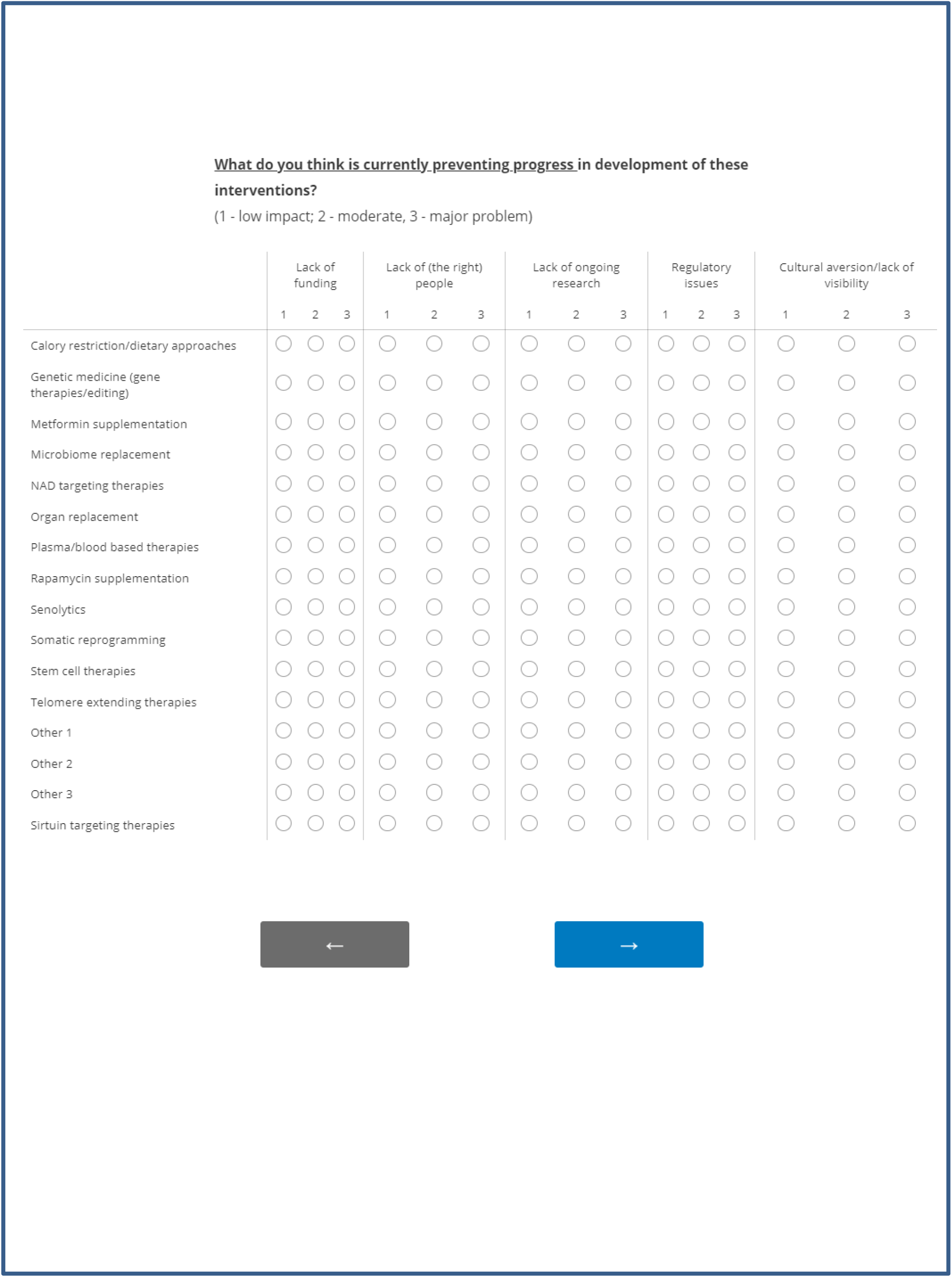
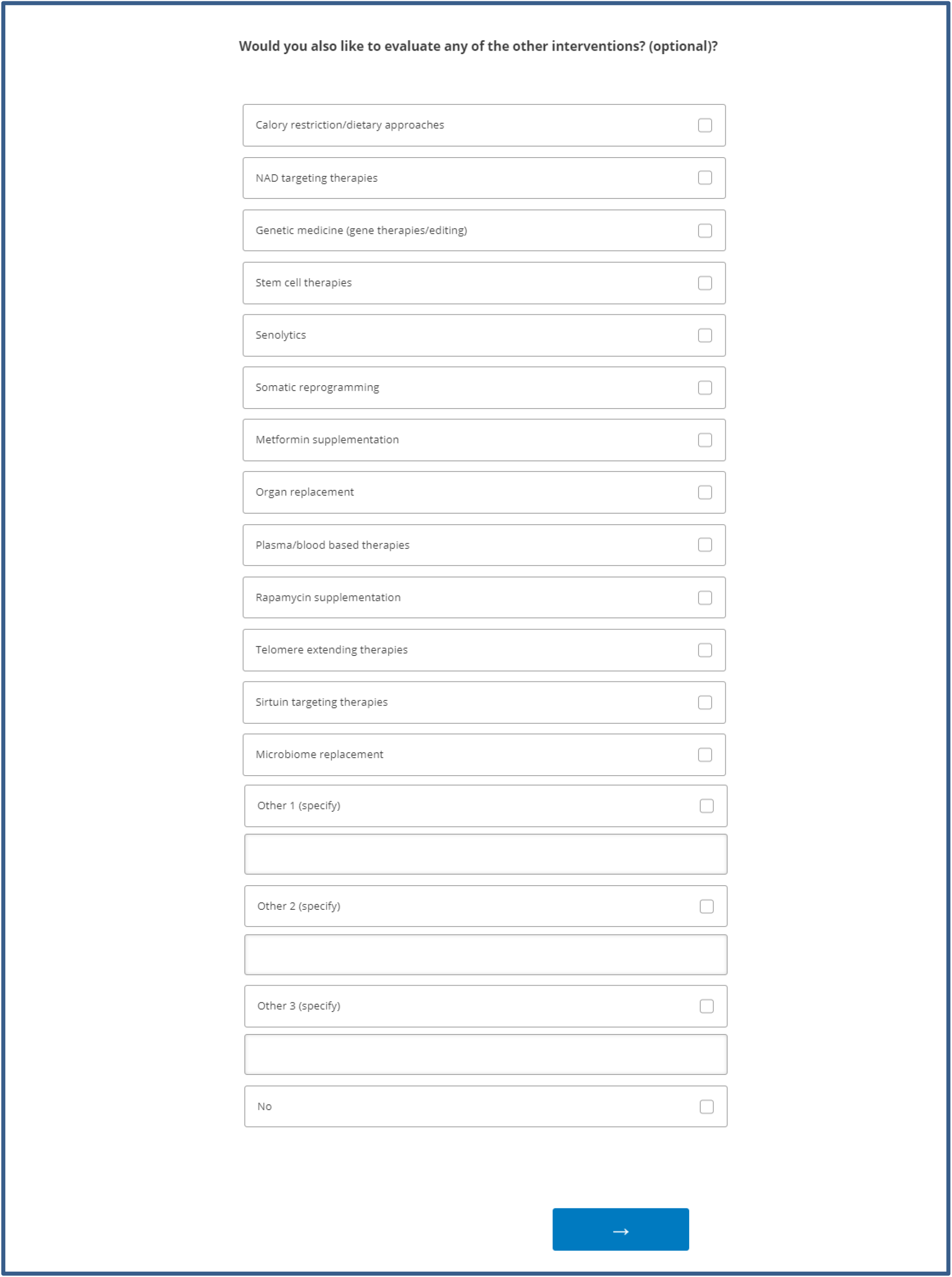
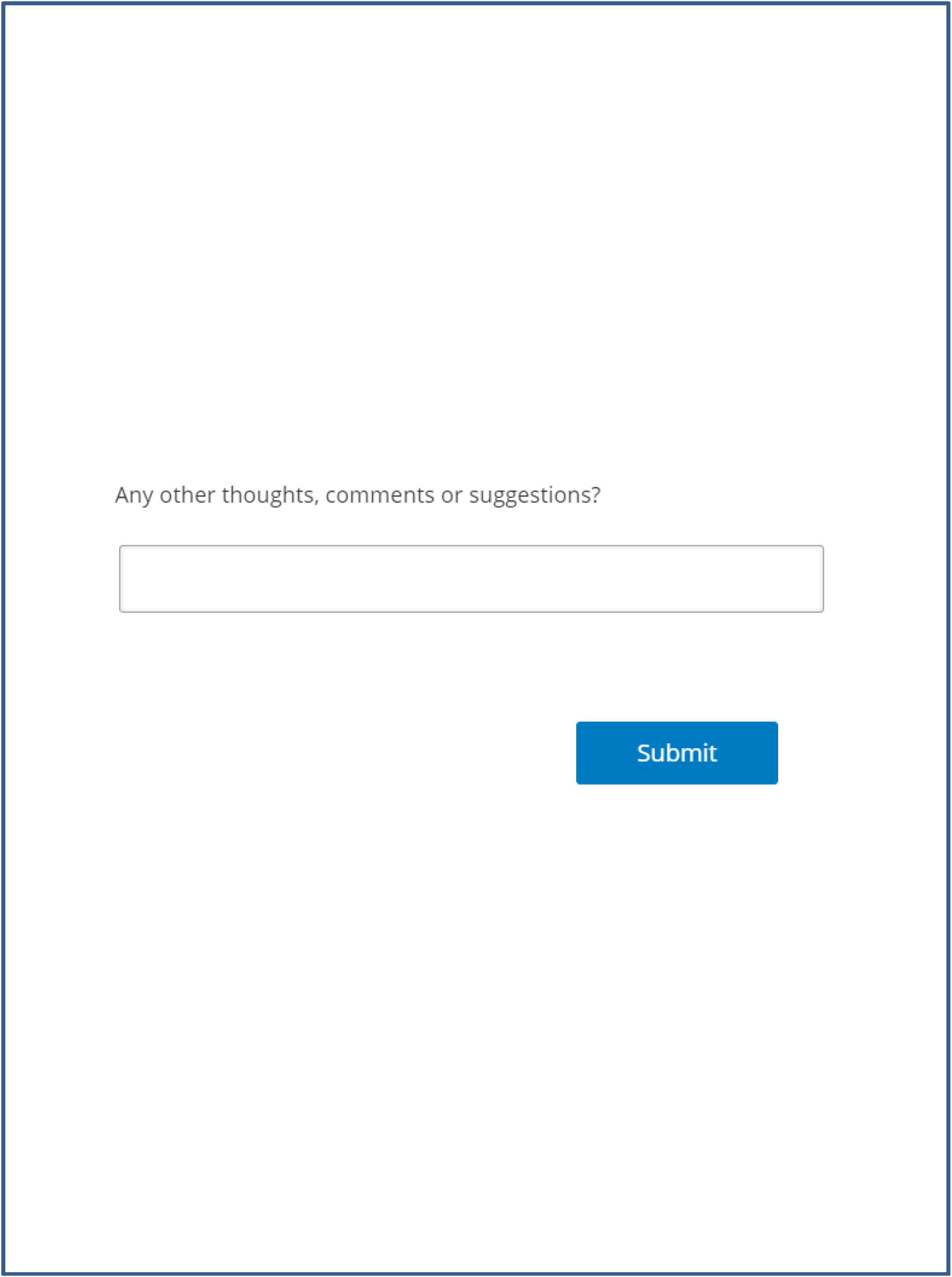
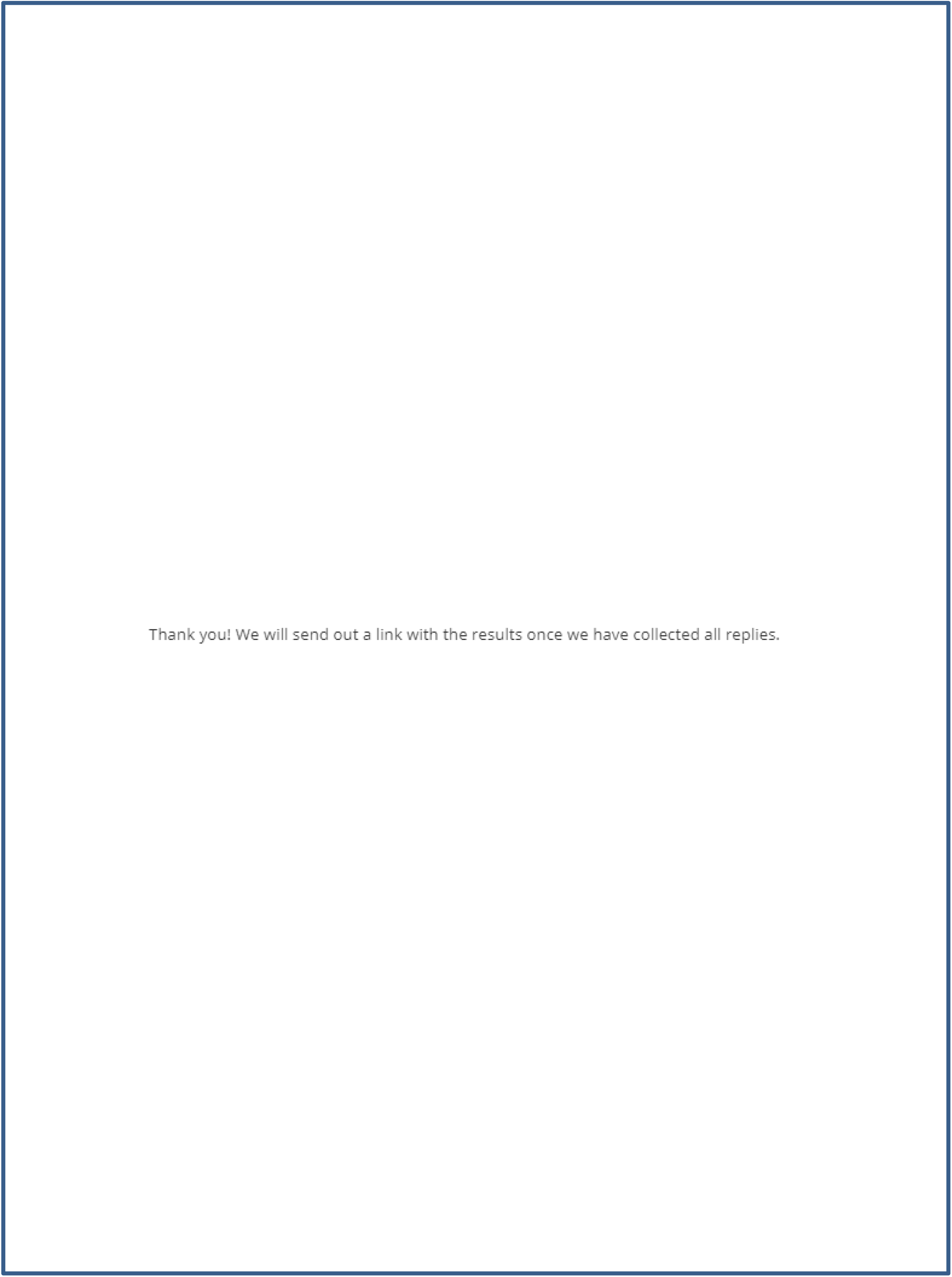
Full survey.

**Figure S2.**
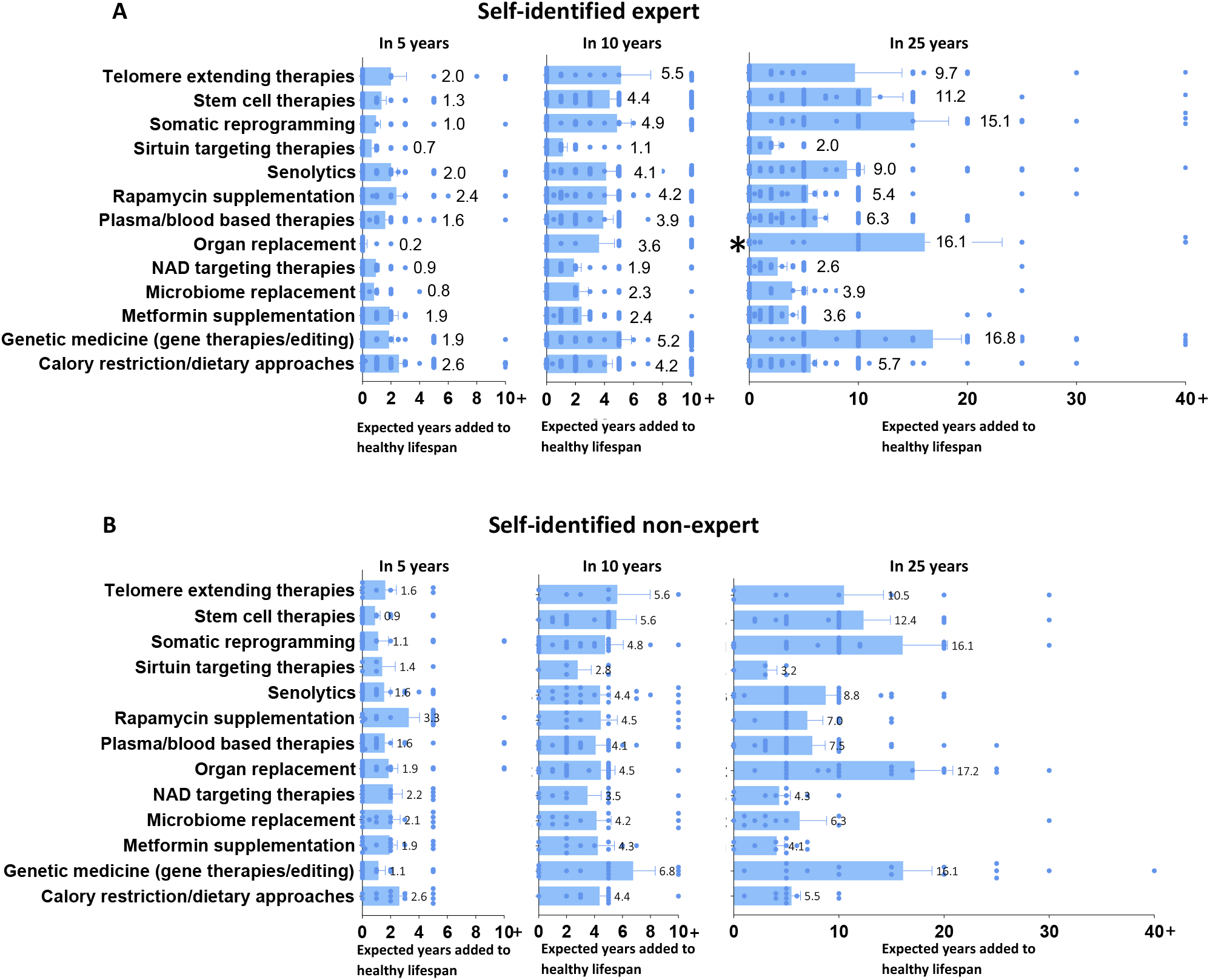
Comparison of self-identified expert vs non-expert estimations of intervention efficacies. **(A)** Self-identified expert evaluations with one (200 year) estimate removed for organ replacement on the right graph (marked with *). **(B)** Self-identified non-expert evaluations. Error bars represent standard error of the mean (SEM).

**Figure S3.**
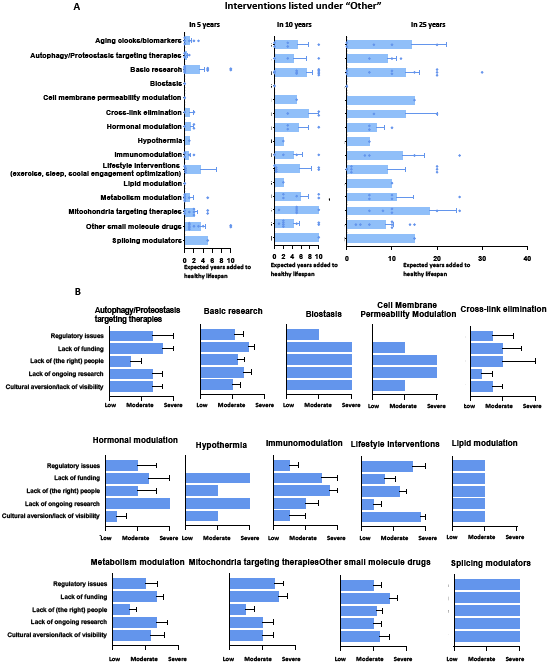
Interventions listed as “Other” by self-identified experts. **(A)** Perceived efficacy of interventions. Individual dots represent individual estimates. **(B)** Bottlenecks for the interventions. Error bars denote standard error of the mean (SEM). In graphs where bars or error bars are missing, this is due to N=1 or missing data (not rated by evaluators).

**Table S1.**
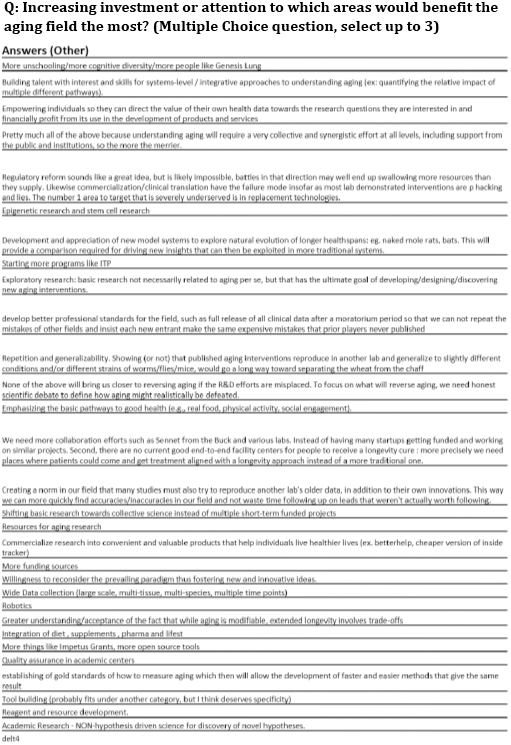
Fig. 2B answers listed as “Other”.

**Table S2.**
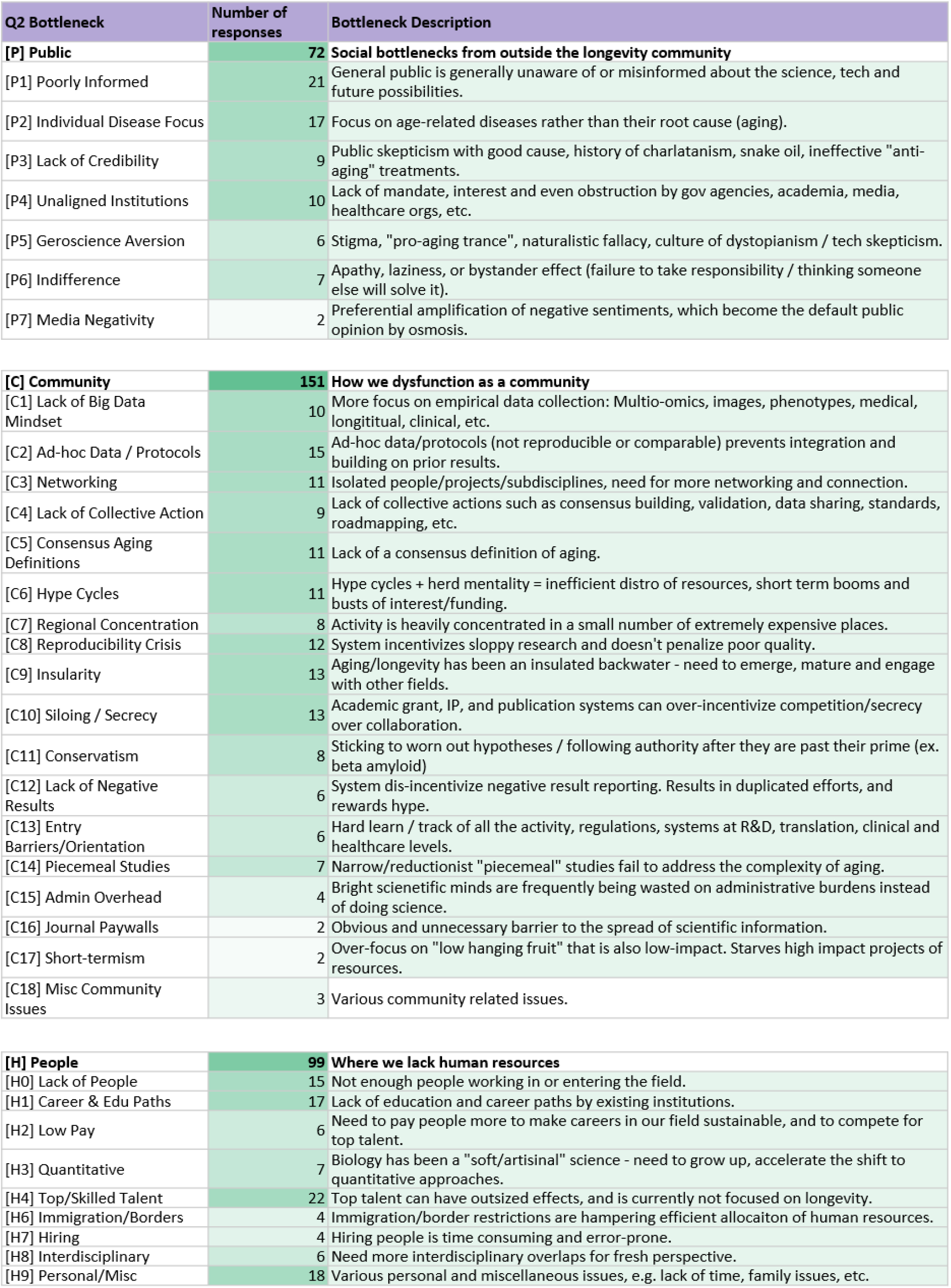
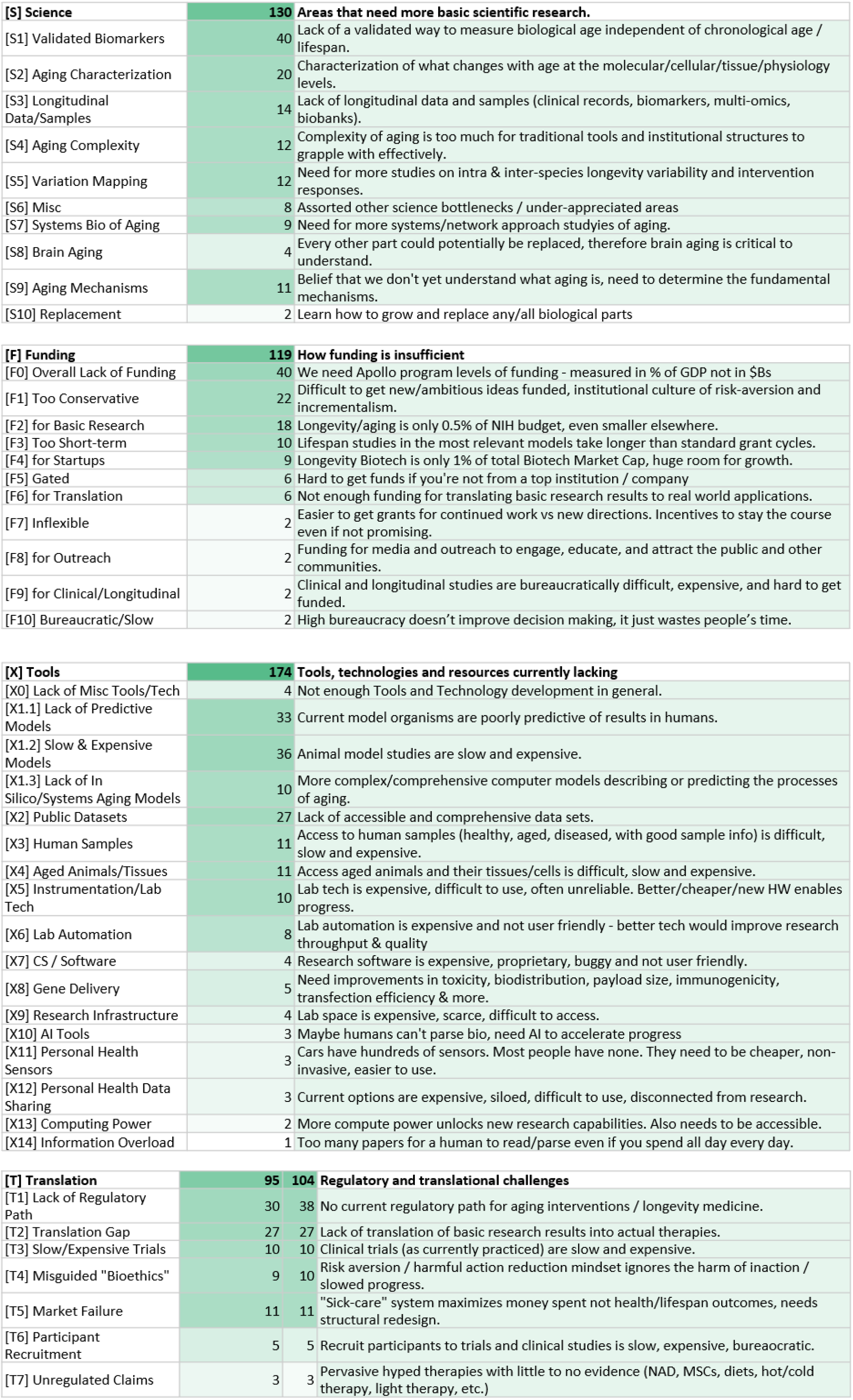
Categories, Bottlenecks and descriptions identified from the survey (described in Fig. 3)

| Q2 Bottleneck | Number of responses | Bottleneck Description |
| --- | --- | --- |
| **[P] Public** | **72** | **Social bottlenecks from outside the longevity community** |
| [P1] Poorly Informed | 21 | General public is generally unaware of or misinformed about the science, tech and future possibilities. |
| [P2] Individual Disease Focus | 17 | Focus on age-related diseases rather than their root cause (aging). |
| [P3] Lack of Credibility | 9 | Public skepticism with good cause, history of charlatanism, snake oil, ineffective "anti-aging" treatments. |
| [P4] Unaligned Institutions | 10 | Lack of mandate, interest and even obstruction by gov agencies, academia, media, healthcare orgs, etc. |
| [P5] Geroscience Aversion | 6 | Stigma, "pro-aging trance", naturalistic fallacy, culture of dystopianism / tech skepticism. |
| [P6] Indifference | 7 | Apathy, laziness, or bystander effect (failure to take responsibility / thinking someone else will solve it). |
| [P7] Media Negativity | 2 | Preferential amplification of negative sentiments, which become the default public opinion by osmosis. |
| **[C] Community** | **151** | **How we dysfunction as a community** |
| [C1] Lack of Big Data Mindset | 10 | More focus on empirical data collection: Multio-omics, images, phenotypes, medical, longitudinal, clinical, etc. |
| [C2] Ad-hoc Data / Protocols | 15 | Ad-hoc data/protocols (not reproducible or comparable) prevents integration and building on prior results. |
| [C3] Networking | 11 | Isolated people/projects/subdisciplines, need for more networking and connection. |
| [C4] Lack of Collective Action | 9 | Lack of collective actions such as consensus building, validation, data sharing, standards, roadmapping, etc. |
| [C5] Consensus Aging Definitions | 11 | Lack of a consensus definition of aging. |
| [C6] Hype Cycles | 11 | Hype cycles + herd mentality = inefficient distro of resources, short term booms and busts of interest/funding. |
| [C7] Regional Concentration | 8 | Activity is heavily concentrated in a small number of extremely expensive places. |
| [C8] Reproducibility Crisis | 12 | System incentivizes sloppy research and doesn't penalize poor quality. |
| [C9] Insularity | 13 | Aging/longevity has been an insulated backwater - need to emerge, mature and engage with other fields. |
| [C10] Siloing / Secrecy | 13 | Academic grant, IP, and publication systems can over-incentivize competition/secrecy over collaboration. |
| [C11] Conservatism | 8 | Sticking to worn out hypotheses / following authority after they are past their prime (ex. beta amyloid) |
| [C12] Lack of Negative Results | 6 | System disincentivize negative result reporting. Results in duplicated efforts, and rewards hype. |
| [C13] Entry Barriers/Orientation | 6 | Hard learn / track of all the activity, regulations, systems at R&D, translation, clinical and healthcare levels. |
| [C14] Piecemeal Studies | 7 | Narrow/reductionist "piecemeal" studies fail to address the complexity of aging. |
| [C15] Admin Overhead | 4 | Bright scientific minds are frequently being wasted on administrative burdens instead of doing science. |
| [C16] Journal Paywalls | 2 | Obvious and unnecessary barrier to the spread of scientific information. |
| [C17] Short-termism | 2 | Over-focus on "low hanging fruit" that is also low-impact. Starves high impact projects of resources. |
| [C18] Misc Community Issues | 3 | Various community related issues. |
| **[H] People** | **99** | **Where we lack human resources** |
| [H0] Lack of People | 15 | Not enough people working in or entering the field. |
| [H1] Career & Edu Paths | 17 | Lack of education and career paths by existing institutions. |
| [H2] Low Pay | 6 | Need to pay people more to make careers in our field sustainable, and to compete for top talent. |
| [H3] Quantitative | 7 | Biology has been a "soft/artisinal" science - need to grow up, accelerate the shift to quantitative approaches. |
| [H4] Top/Skilled Talent | 22 | Top talent can have outsized effects, and is currently not focused on longevity. |
| [H6] Immigration/Borders | 4 | Immigration/border restrictions are hampering efficient allocation of human resources. |
| [H7] Hiring | 4 | Hiring people is time consuming and error-prone. |
| [H8] Interdisciplinary | 6 | Need more interdisciplinary overlaps for fresh perspective. |
| [H9] Personal/Misc | 18 | Various personal and miscellaneous issues, e.g. lack of time, family issues, etc. |
| **[S] Science** | **130** | **Areas that need more basic scientific research.** |
| [S1] Validated Biomarkers | 40 | Lack of a validated way to measure biological age independent of chronological age / lifespan. |
| [S2] Aging Characterization | 20 | Characterization of what changes with age at the molecular/cellular/tissue/physiology levels. |
| [S3] Longitudinal Data/Samples | 14 | Lack of longitudinal data and samples (clinical records, biomarkers, multi-omics, biobanks). |
| [S4] Aging Complexity | 12 | Complexity of aging is too much for traditional tools and institutional structures to grapple with effectively. |
| [S5] Variation Mapping | 12 | Need for more studies on intra & inter-species longevity variability and intervention responses. |
| [S6] Misc | 8 | Assorted other science bottlenecks / under-appreciated areas |
| [S7] Systems Bio of Aging | 9 | Need for more systems/network approach studies of aging. |
| [S8] Brain Aging | 4 | Every other part could potentially be replaced, therefore brain aging is critical to understand. |
| [S9] Aging Mechanisms | 11 | Belief that we don't yet understand what aging is, need to determine the fundamental mechanisms. |
| [S10] Replacement | 2 | Learn how to grow and replace any/all biological parts |
| **[F] Funding** | **119** | **How funding is insufficient** |
| [F0] Overall Lack of Funding | 40 | We need Apollo program levels of funding - measured in % of GDP not in \$Bs |
| [F1] Too Conservative | 22 | Difficult to get new/ambitious ideas funded, institutional culture of risk-aversion and incrementalism. |
| [F2] for Basic Research | 18 | Longevity/aging is only 0.5% of NIH budget, even smaller elsewhere. |
| [F3] Too Short-term | 10 | Lifespan studies in the most relevant models take longer than standard grant cycles. |
| [F4] for Startups | 9 | Longevity Biotech is only 1% of total Biotech Market Cap, huge room for growth. |
| [F5] Gated | 6 | Hard to get funds if you're not from a top institution / company |
| [F6] for Translation | 6 | Not enough funding for translating basic research results to real world applications. |
| [F7] Inflexible | 2 | Easier to get grants for continued work vs new directions. Incentives to stay the course even if not promising. |
| [F8] for Outreach | 2 | Funding for media and outreach to engage, educate, and attract the public and other communities. |
| [F9] for Clinical/Longitudinal | 2 | Clinical and longitudinal studies are bureaucratically difficult, expensive, and hard to get funded. |
| [F10] Bureaucratic/Slow | 2 | High bureaucracy doesn't improve decision making, it just wastes people's time. |
| **[X] Tools** | **174** | **Tools, technologies and resources currently lacking** |
| [X0] Lack of Misc Tools/Tech | 4 | Not enough Tools and Technology development in general. |
| [X1.1] Lack of Predictive Models | 33 | Current model organisms are poorly predictive of results in humans. |
| [X1.2] Slow & Expensive Models | 36 | Animal model studies are slow and expensive. |
| [X1.3] Lack of In Silico/Systems Aging Models | 10 | More complex/comprehensive computer models describing or predicting the processes of aging. |
| [X2] Public Datasets | 27 | Lack of accessible and comprehensive data sets. |
| [X3] Human Samples | 11 | Access to human samples (healthy, aged, diseased, with good sample info) is difficult, slow and expensive. |
| [X4] Aged Animals/Tissues | 11 | Access aged animals and their tissues/cells is difficult, slow and expensive. |
| [X5] Instrumentation/Lab Tech | 10 | Lab tech is expensive, difficult to use, often unreliable. Better/cheaper/new HW enables progress. |
| [X6] Lab Automation | 8 | Lab automation is expensive and not user friendly - better tech would improve research throughput & quality |
| [X7] CS / Software | 4 | Research software is expensive, proprietary, buggy and not user friendly. |
| [X8] Gene Delivery | 5 | Need improvements in toxicity, biodistribution, payload size, immunogenicity, transfection efficiency & more. |
| [X9] Research Infrastructure | 4 | Lab space is expensive, scarce, difficult to access. |
| [X10] AI Tools | 3 | Maybe humans can't parse bio, need AI to accelerate progress |
| [X11] Personal Health Sensors | 3 | Cars have hundreds of sensors. Most people have none. They need to be cheaper, non-invasive, easier to use. |
| [X12] Personal Health Data Sharing | 3 | Current options are expensive, siloed, difficult to use, disconnected from research. |
| [X13] Computing Power | 2 | More compute power unlocks new research capabilities. Also needs to be accessible. |
| [X14] Information Overload | 1 | Too many papers for a human to read/parse even if you spend all day every day. |

|  |  |  |  |
| --- | --- | --- | --- |
| **[T] Translation** | **95** | **104** | **Regulatory and translational challenges** |
| [T1] Lack of Regulatory Path | 30 | 38 | No current regulatory path for aging interventions / longevity medicine. |
| [T2] Translation Gap | 27 | 27 | Lack of translation of basic research results into actual therapies. |
| [T3] Slow/Expensive Trials | 10 | 10 | Clinical trials (as currently practiced) are slow and expensive. |
| [T4] Misguided "Bioethics" | 9 | 10 | Risk aversion / harmful action reduction mindset ignores the harm of inaction / slowed progress. |
| [T5] Market Failure | 11 | 11 | "Sick-care" system maximizes money spent not health/lifespan outcomes, needs structural redesign. |
| [T6] Participant Recruitment | 5 | 5 | Recruit participants to trials and clinical studies is slow, expensive, bureaucratic. |
| [T7] Unregulated Claims | 3 | 3 | Pervasive hyped therapies with little to no evidence (NAD, MSCs, diets, hot/cold therapy, light therapy, etc.) |

**Table S3.** Solutions Categories, tags and descriptions identified from the survey (described in Fig. 4)

| Q3 Solution | Answers | Solution Description |
| --- | --- | --- |
| **[+P] Public** | **30** | **Social projects aimed outside of the longevity community** |
| [+P1.1] Outreach to Public | 11 | Shift public sentiment towards pro-longevity. Cascading benefits (policy, funding, market, career choice...) |
| [+P1.2] Outreach to Other Fields | 2 | Bring fresh perspectives and skillsets into the longevity movement. |
| [+P1.3] Outreach to Biotech/Pharma | 1 | Only 1% of Biotech/Pharma is longevity/aging oriented, lots of room for growth. |
| [+P1.4] Outreach to Students | 0 | Deflect career paths at the highest leverage point. |
| [+P1.5] Outreach to Gov/Institutions | 3 | Engage/infiltrate to improve alignment. They have the power to break many bottlenecks if aligned. |
| [+P2] Culture | 10 | A widespread pro-longevity cultural movement. |
| [+P3] Incentives | 2 | Regulations/policies to incentivize longevity practices/industry growth - works for climate why not longevity? |
| [+P4] Comms Training | 1 | For scientists, engineers, operators in longevity - training to communicate clearly and effectively to the public. |
| **[+C] Community** | **73** | **How we could improve functioning of the community** |
| [+C1] Big Data Mindset | 2 | More focus on empirical data collection: Multi-omics, images, phenotypes, medical, longitudinal, clinical, etc. |
| [+C2] Data/Protocol Standards | 11 | Set and update standards for data and protocols, incentivize their use. |
| [+C3] Networking | 13 | Mechanism to connect employers, entrepreneurs, investors, volunteers, etc. |
| [+C4] Large Collaborative Projects | 8 | Large collaborative projects and rather than ego/grant/publication driven competition. |
| [+C5] Collective Action | 8 | Organize to facilitate consensus building, validation, data sharing, standards, roadmapping, etc. |
| [+C6] Consensus Aging Definition | 6 | Reach a consensus theory/definition of aging |
| [+C7] New Research Centers | 6 | More dedicated aging/longevity research centers. Diverse locations. Varied specialties. Facilitate large projects. |
| [+C8] Reproduce Key Results | 3 | Funded centers dedicated to reproducing / testing the robustness of key results. |
| [+C9] Field-Wide Status Dashboard | 7 | Overview of all proposals, projects, pathways, targets, pre and in-clinic pipelines. Linked to data. |
| [+C10] Outsourcing Services | 4 | Longevity CROs/Cores - make outsourcing research easier such as lifespan studies, multi-omic analysis. |
| [+C11] Neg Results Incentives | 3 | Mechanism to incentivize/enable reporting negative results. Pre-results publication approvals for example. |
| [+C12] Open Science | 1 | Make data, samples, results open and available by default. |
| [+C13] Long-termism | 1 | More focus on long term research and technology paths with higher-impact potential (vs "low hanging fruit") |
| [+C14] Protected Time | 1 | Reduce/limit the amount of time spent on non-research activities (admin, grants, etc.) |
| [+C15] Leadership | 1 | Bold and ambitious, yet free of ego/hype and relatable. Leadership is lacking but necessary to build a movement. |
| **[+S] Science** | **35** | **Research goals, that if achieved would unlock progress.** |
| [+S1] Validated Biomarkers | 13 | Validated way to measure biological age independent of chronological age / lifespan. |
| [+S2] Aging Fully Characterized | 10 | Comprehensive characterization of aging at the molecular/cellular/tissue/physiology levels. |
| [+S3] Longitudinal Data/Samples | 8 | Unlock or generate more longitudinal data and samples (clinical records, biomarkers, multi-omics, biobanks) |
| [+S4] Aging Variation Mapped | 3 | Comprehensive map of Intra and inter species aging variation (omics, physiology, phenotype, etc.) |
| [+S10] Brain Rejuvenation/Replacement | 1 | Ambitious and broad range of R&D on brain rejuvenation/replacement. |
| [+S11] Comprehensive Replacement | 0 | Methods to grow and replace any/all biological parts |

**Table S4.**

|  |  |  |
| --- | --- | --- |
| **[+F] Funding** | **47** | **How to improve funding** |
| [+F0] Overall More Funding | 18 | We need Apollo program levels of funding - measured in % of GDP not in \$Bs |
| [+F1] Ambitious/Exploratory | 11 | More ambitious/exploratory projects are needed to accelerate progress. |
| [+F2] for Basic Research | 5 | Longevity/aging is only 0.5% of NIH budget, even smaller elsewhere. |
| [+F3] Longer-term | 6 | Longer bets on people and projects are necessary for many aging research directions. |
| [+F4] for Startups | 3 | Longevity Biotech is only 1% of total Biotech Market Cap, huge room for growth. |
| [+F5] for Outsiders | 1 | Take more bets on system outsiders (non-academic / non-ivy league). |
| [+F6] for Translation | 1 | Not enough funding for translating basic research results to real world applications. |
| [+F7] Flexible | 1 | Grants for promising people to transition from other fields/areas. |
| [+F8] for Outreach | 0 | Funding for media and outreach to engage, educate, and attract the public and other communities. |
| [+F9] for Clinical/Longitudinal | 0 | Clinical and longitudinal studies are bureaucratically difficult, expensive, and hard to get funded. |
| [+F10] Faster Grants | 1 | High bureaucracy doesn't improve decision making, it just wastes people's time. |
| **[+H] People** | **18** | **Solutions for human resources** |
| [+H0] More People | 2 | Not enough people working in or entering the field. |
| [+H1] Quantitative | 1 | Need to recruit or train more people in quantitative skills (math, physics, statistics, ml, comp/systems bio) |
| [+H2] Higher Pay | 3 | Need to pay people more to make careers in our field sustainable, and to compete for top talent. |
| [+H3] Longevity Training | 2 | We need edu/career paths for people to start or shift careers in longevity biotech. |
| [+H4] Skills Assessment | 2 | Better alternatives to degrees/resumes/interviews? Get people to work faster and more efficiently. |
| [+H5] Interdisciplinary | 1 | Need more interdisciplinary overlaps for fresh perspective. |
| [+H6] More/Flexible Visas | 1 | Immigration/border restrictions are hampering efficient allocation of human resources. |
| [+H10] Personal/Misc | 4 | Misc personal solutions |
| [+H9] Top/Skilled Talent | 2 | Top talent can have outsized effects, and is currently not focused on longevity. |
| **[+X] Tools** | **79** | **Needed tools, technologies and resources** |
| [+X0] Unspecified Tools/Tech | 1 | More Tools and Technology development in general. |
| [+X1.1] In Silico/Systems Aging Models | 4 | Computational/Systems Bio Modeling. Make predictions in silico to save time and cost in meatspace. |
| [+X1.2] Ex vivo Models | 2 | Cell/tissue/organ based aging models that are predictive, affordable and accessible. |
| [+X1.3] Multiple/Diverse Models | 3 | Improve robustness of results by using multiple models. |
| [+X1.4] Unspecified/Misc Models | 3 | Better model systems for aging (unspecified how) |
| [+X1.5] Longer Lived Models | 2 | Develop longer lived animal models - more likely to be more translatable to human biology. |
| [+X1.6] Human Models | 2 | Skip animals and their problems - develop human model systems (cell, tissue, organ, body, in vivo) |
| [+X1.7] Accelerated Aging Models | 1 | Accelerated aging models could speed progress... if they are representative of normal aging. |
| [+X2] Public Datasets | 22 | Accessible and comprehensive data sets can lift all boats, accelerate progress. |
| [+X3] Instrumentation/Lab Tech | 8 | Cheaper/better/new tools to measure biology/health/age would enable more and new studies. |
| [+X4] Human Cell/Tissue Bank | 6 | Easy, inexpensive, and well documented samples. |
| [+X5] Aged Animals/Tissue Bank | 5 | Easy, inexpensive, and well documented aged animals and samples. |
| [+X5] Lab Automation | 4 | Better/easier lab automation would enable increased throughput, higher quality, standards, collaboration. |
| [+X6] CS / Software | 5 | Better/easier research software would enable increased throughput, higher quality, standards, collaborations. |
| [+X7] Gene Delivery | 1 | Better delivery and gene editing tools |
| [+X8] AI Tools | 4 | Overcome the limits of human intelligence by building better/specialized AI systems. |
| [+X9] Personal Health Sensors | 2 | Cars have hundreds of sensors. Most people have none. They need to be cheaper, non-invasive, easier to use. |
| [+X10] Syn Bio | 0 | Tools to engineer cells, tissues, organs to meet our specifications. |
| [+X11] Computing Power | 1 | More compute power unlocks new research capabilities. Also needs to be accessible. |
| [+X12] Health Data Platform | 3 | Platform for individuals to easily analyze and share health and medical data for advancing research. |
| **[+T] Translation** | **33** | **How to improve translational challenges** |
| [+T1] Regulatory Path | 19 | A regulatory path for aging interventions / longevity medicine. |
| [+T2] Participant Database | 4 | Database/marketplace to streamline sourcing clinical research participants. |
| [+T3] Regulatory Opt-Out | 5 | Allow people to opt-out of medical regulations. A healthcare socioeconomic free zone. |
| [+T4] Intervention Testing | 0 | More intervention testing, including combination therapies. |
| [+T5] Translation Bridges | 2 | Innovation centers and funding to close the translation gap. |
| [+T6] Regulate Claims | 1 | Require trials to make health claims in supplements, lifestyle, etc. Opportunity to design/test lower cost trials? |
| [+T7] Public Trials | 0 | No current mechanism to test interventions when a therapy isn't IP protectable / highly profitable |
| [+T8] Rethink Trials | 1 | How can we make them bigger/better/cheaper and longevity oriented. |
| [+T8] Data Unlocking | 1 | Change policies to unlock siloed health/medical data. Require accessibility, standards and interoperability. |
| [+T9] Longevity Standard of Care | 0 | Alternative to "sick-care" standards, to help create a market for longevity medicine. |

